# Selective Enhancer Dependencies in *MYC*-Intact and *MYC*-Rearranged Germinal Center B-cell Diffuse Large B-cell Lymphoma

**DOI:** 10.1101/2023.05.02.538892

**Authors:** Ashwin R. Iyer, Aishwarya Gurumurthy, Rohan Kodgule, Athalee R. Aguilar, Travis Saari, Abdullah Ramzan, Dylan Rausch, Juhi Gupta, Cody N. Hall, John S. Runge, Matthew Weiss, Mahshid Rahmat, Rockwell Anyoha, Charles P. Fulco, Irene M. Ghobrial, Jesse Engreitz, Marcin P. Cieslik, Russell J.H. Ryan

## Abstract

High expression of *MYC* and its target genes define a subset of germinal center B-cell diffuse large B-cell lymphoma (GCB-DLBCL) associated with poor outcomes. Half of these high-grade cases show chromosomal rearrangements between the *MYC* locus and heterologous enhancer-bearing loci, while focal deletions of the adjacent non-coding gene *PVT1* are enriched in *MYC*-intact cases. To identify genomic drivers of *MYC* activation, we used high-throughput CRISPR-interference (CRISPRi) profiling of candidate enhancers in the *MYC* locus and rearrangement partner loci in GCB-DLBCL cell lines and mantle cell lymphoma (MCL) comparators that lacked common rearrangements between *MYC* and immunoglobulin (Ig) loci. Rearrangements between *MYC* and non-Ig loci were associated with unique dependencies on specific enhancer subunits within those partner loci. Notably, fitness dependency on enhancer modules within the *BCL6* super-enhancer (*BCL6*-SE) cluster regulated by a transcription factor complex of MEF2B, POU2F2, and POU2AF1 was higher in cell lines bearing a recurrent *MYC::BCL6*-SE rearrangement. In contrast, GCB-DLBCL cell lines without *MYC* rearrangement were highly dependent on a previously uncharacterized 3’ enhancer within the *MYC* locus itself (GCBME-1), that is regulated in part by the same triad of factors. GCBME-1 is evolutionarily conserved and active in normal germinal center B cells in humans and mice, suggesting a key role in normal germinal center B cell biology. Finally, we show that the *PVT1* promoter limits *MYC* activation by either native or heterologous enhancers and demonstrate that this limitation is bypassed by 3’ rearrangements that remove *PVT1* from its position in *cis* with the rearranged *MYC* gene.

**Key points:**

- CRISPR-interference screens identify a conserved germinal center B cell *MYC* enhancer that is essential for GCB-DLBCL lacking *MYC* rearrangements.
- Functional profiling of *MYC* partner loci reveals principles of *MYC* enhancer-hijacking activation by non-immunoglobulin rearrangements.

## Introduction

Diffuse large B-cell lymphoma (DLBCL) is a clinically heterogeneous cancer for which distinctive biological subgroups have been defined ^1–3^. The most aggressive subgroup known as “double-hit signature” (DHITsig+)^4^ or “molecular high-grade” (MHG)^5^ is characterized by a germinal center B cell phenotype and high expression of *MYC* and *MYC* target genes. The best-characterized mechanisms of *MYC* hyperactivity in DLBCL are long distance rearrangements that juxtapose *MYC* to well-studied immunoglobulin (Ig) locus enhancers^6, 7^, driving *MYC* expression above physiological levels^8^. *MYC* rearrangements to non-Ig loci are assumed to activate *MYC* by a similar “enhancer hijacking” mechanism, but our functional understanding of these events is limited. Even less is known about how *MYC* is activated in the roughly half of DHITsig/MHG-DLBCL that lack detectable *MYC* locus rearrangements by fluorescence in situ hybridization (FISH)^4, 9^ or cryptic insertions of heterologous enhancers into the *MYC* locus as identified by whole-genome-sequencing in a small proportion of additional cases^9^. To address these questions, we performed high-throughput CRISPR-interference (CRISPRi) screens targeting candidate enhancers in the *MYC* locus and rearrangement partner loci, revealing regulatory principles of *MYC*-non-Ig rearrangements, and showing that a previously uncharacterized enhancer active in normal germinal center B cells serves as a common dependency of *MYC*-intact GCB-DLBCL.

## Materials and Methods

### Cell Lines and lentivirus production

See Supplementary Table 1 for details of cell line source, validation and media. VSV-G - pseudotyped lentivirus was generated in 293T cells by standard methods.

### ChIP-Seq and ATAC-Seq

H3K27ac ChIP-Seq was performed as previously described^10^, with formaldehyde crosslinking, nuclear isolation, sonication in 0.3% SDS lysis buffer with a Q800R2 Sonifier (QSonica), dilution to 0.1% SDS, and rotation at 4°C overnight with 2 ug of antibody (H3K27ac, #39133, Active Motif). Transcription factor ChIP-Seq was performed as previously described^11^ with 2-5 ug of antibody was used per immunoprecipitation (POU2F2, sc-233X, Santa Cruz; POU2AF1, sc-955X, Santa Cruz; p300, A300-358A, Bethyl). ATAC-Seq was performed as previously described^10^ with nuclei isolated from 50,000 cells for each sample. See Supplementary Methods for details.

### CRISPRi assays

Doxycycline-inducible KRAB-dCas9-expressing cell lines were generated as previously described^12^. Briefly, GCB-DLBCL and MCL cell lines were co-transduced with lentivirus produced from TRE-KRAB-dCas9-IRES-GFP (Addgene #85556) and EF1a_TetOn3G (Clontech) and serially sorted to derive populations that were uniformly GFP-negative before and GFP-positive after doxycycline induction. Cell lines were validated by showing differential loss of viability when transduced with sgRNAs targeting essential genes (RPL34 and RPL8) versus non-targeting controls upon doxycycline induction. For the cell line DB, we derived cells stably transduced with a constitutive KRAB-dCas9 vector, UCOE-SFFV-KRAB-dCas9-P2A-mCherry, generated through ligation of the promoter region from pMH001 (Addgene #85969) in place of the promoter of pHR-SFFV-KRAB-dCas9-P2A-mCherry (Addgene #60954) following AgeI and MluI digestion.

For single sgRNA studies, 20-mer seed sequences with a prepended “G” were restriction cloned into BsmBI-digested sgOpti (Addgene #85681). Inducible dCas9-KRAB expressing cells were transduced in triplicate with sgOpti lentivirus and divided at 24 hours into media with 1 ug/mL puromycin with or without 500 ng / mL doxycycline. Viable cellularity was quantified at day 7 with CellTiter-Glo reagent (Promega) according to manufacturer’s instructions. For gene expression studies, 1 million cells per well were transduced with sgRNA lentivirus, transferred at 24 hours to media with 1 ng/ml puromycin and 500 ng/ml doxycycline, and harvested 3 or 5 days after transduction for RNA extraction and qRT-PCR. For dual sgRNA studies, sgRNA seed sequences were cloned into derivatives of sgOpti bearing tagBFP (pMW-tagBFP) or tagRFP (pMW-tagRFP) reporters. Inducible KRAB-dCas9-bearing cell lines were transduced with two sgRNA lentiviral supernatants bearing different reporters, sorted for cells with dual reporter expression, and were then induced with 100-500 ng/mL doxycycline for three days (gene expression) or 7 days (viable cellularity).

### High-throughput CRISPRi screening

Tiling screens covering candidate *MYC* enhancer regions were conducted with a published sgRNA library^12^ designed to cover broad regions of the *MYC* locus. For the NFR-focused sgRNA library targeting the *MYC* and rearrangement partner loci, 500 bp-intervals centered on ATAC-Seq peaks and associated with a high H3K27ac / ATAC-Seq activity score^13^ in at least one of 26 mature B-cell lymphoma cell lines were merged across cell lines. FlashFry^14^ was used to identify and score sgRNAs within these intervals, which were filtered to retain sgRNAs with low predicted off-target and high predicted on-target effects, and downsampled to 35 sgRNA per 1 kb. sgRNA library cloning was performed as described^15^. CRISPRi screening was performed in doxycycline-inducible KRAB-dCas9 expressing cell lines as previously described, with 2-3 replicates per tiling screen and 3-4 replicates per NFR-focused screen. For the DB screen conducted in a cell line with non-inducible dCas9-KRAB, the same protocol was followed except that equal numbers of the parental cell line were transduced in parallel and harvested following selection as “T0” replicates. Tiling sgRNA screens were analyzed as previously described^12^. NFR-focused screen analysis was performed with MAGeCK-RRA software^16^, applied separately to NFR-targeting and gene-targeting sgRNA sets. The ‘mageck test’ option ‘–norm-method control’ was used to normalize log2fold changes to the non-targeting control sgRNAs. See **Supplementary text** for additional details of CRISPRi library design and experimental procedures.

### Transcriptional reporters

Enhancer sequences were PCR amplified from genomic DNA, cloned by Gibson assembly into the STARR-seq firefly luciferase validation vector ORI_empty (Addgene: 99297). Vectors ORI_SV40 (Addgene:99309) and ORI_neg.cont (Addgene: 99315) were used as positive and negative controls. Site-directed mutagenesis (Q5 site-directed mutagenesis kit E0554S; NEB) was used to delete 7 nucleotides (‘TTTGCAT’) corresponding to the indicated OCT2 binding motif from NFR 195B. Cell lines were electroporated (Neon, Thermo Scientific) with luciferase and Renilla control (pRL-SV40) plasmids in 1:10 ratio. Two replicate electroporations were each divided into three wells for 48h of culture and readout using the Dual-Glo Luciferase kit (Promega; E2940) in a Bio-Tek Synergy HT Microplate Reader.

### Structural variant analysis

4C-Seq was performed and analyzed as previously described^17–19^. Fresh-frozen cell line pellets were sent to Bionano Genomics for optical mapping. Structural variant read maps were visualized with BioNano Access software v1.7.2. See **Supplementary text** for additional details.

### Western Blotting

Western blot analyses were performed by standard protocols with the following antibodies: MEF2B (HPA004734, Atlas), EBF (sc-137065, Santa Cruz Biotechnology), E2A (sc-349X, Santa Cruz Biotechnology), Bob1 (sc-955X, Santa Cruz Biotechnology), Oct-2 (sc-233X, Santa Cruz Biotechnology), Bcl-6 (sc-7388, Santa Cruz Biotechnology), c-Myc (5605S, Cell Signaling Technology) and CTCF (3418S, Cell Signaling Technology).

## Results

### Functional mapping of native and “hijacked” MYC-activating enhancers

We identified DLBCL cell lines with known *MYC* locus rearrangements to non-immunoglobulin loci^11, 20, 21^ or with no evidence of *MYC* locus aberrations (**Supplementary Table S1**), and defined enhancers with high predicated activity based on H3K27ac ChIP-Seq and ATAC-Seq in the MYC locus and rearrangement partner loci specific to each cell line (**Figure 1A** **and Supplementary Table S2**). We designed a custom sgRNA library targeting the nucleosome-free regions (NFRs) of candidate enhancers, as well as promoter-targeting sgRNAs^22^ for transcription factor genes expressed in B cells, along with positive and negative essentiality controls and extended coverage of the *MYC* and *BCL6* promoter regions.

**Figure 1.**
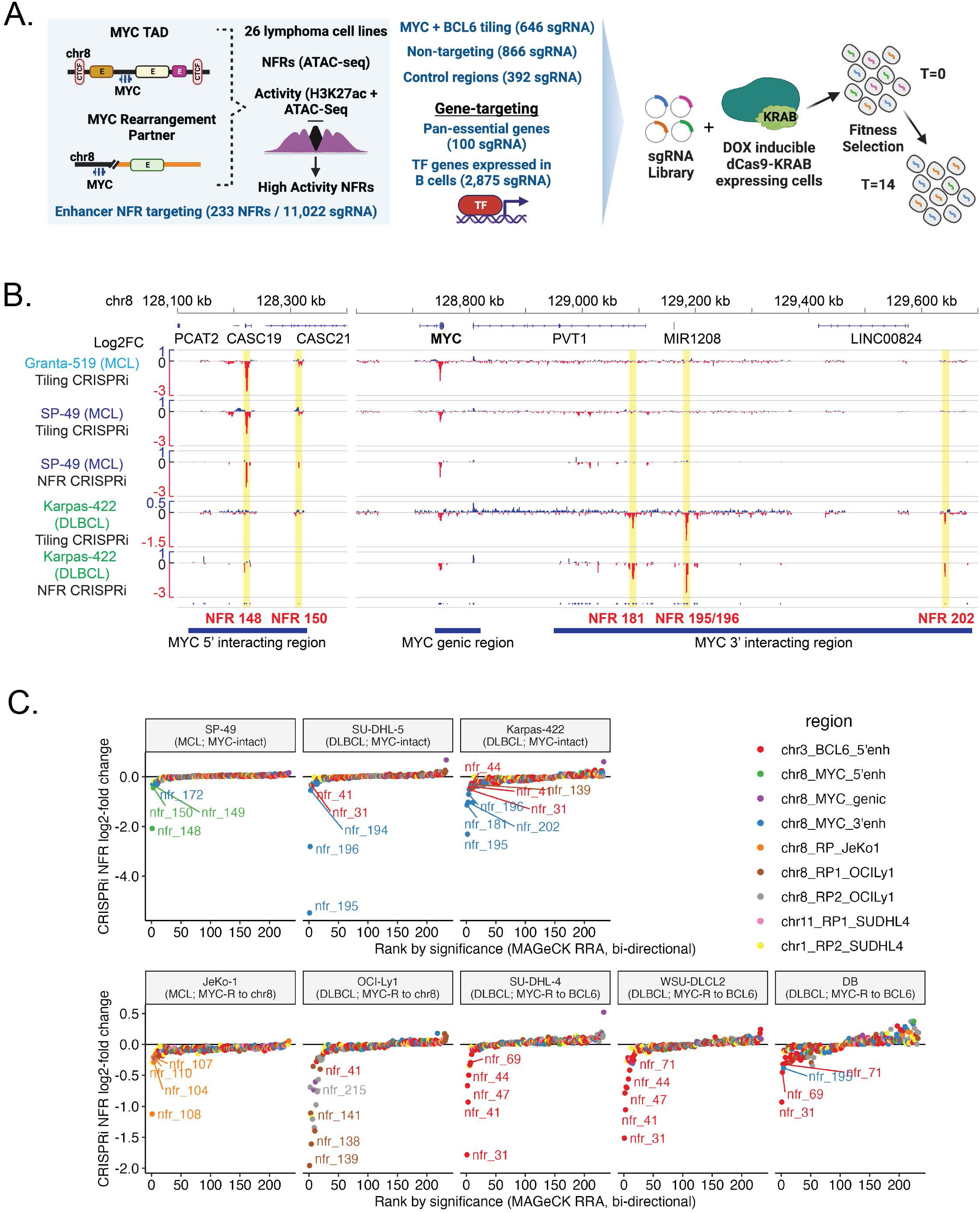
Overview of CRISPRi screening in lymphoma cell lines. **A.** Design strategy for sgRNA library and CRISPRi screen (NFR CRISPRi) targeting candidate enhancers in the *MYC* locus and rearrangement partner regions, transcription factor genes, and controls. **B.** Comparison of CRISPRi screening results in the *MYC* locus with the tiling sgRNA library (y-axis shows log2-fold change averaged over a 20 sgRNA sliding window) and NFR-focused sgRNA library (y-axis shows shows log2-fold of individual sgRNAs) in mantle cell lymphoma (Granta-519, SP-49) and GCB-DLBCL (Karpas-422) cell lines. **C.** Plot of sgRNA depletion or enrichment (MAGeCK RRA log2-fold-change) for targeted intervals in eight lymphoma NFR CRISPRi screens. Target intervals are ranked along the X axis by lowest negative selection FDR p-value (left to right, intervals with log2-fold change <0) or lowest positive selection FDR p-value (right to left, intervals with log2-fold change >0). Intervals are color-coded by genomic region. See B for position of *MYC* target regions. *MYC* rearrangement partner (RP) target regions other than *BCL6* are named for the cell line in which the region is rearranged to *MYC*. See Supplementary Table 2 for positions of all target intervals.

We selected lymphoma cell lines without *MYC* rearrangements for initial CRISPRi screening with both a published *MYC* locus tiling sgRNA library^12^ and our more efficient NFR-focused library targeting both *MYC* and rearrangement partner loci. One NFR and two tiling CRISPRi screens in two MCL cell lines confirmed dependency on 5’ Notch / RBPJ-activated *MYC* enhancers^19, 23^ (**Figure 1B, Supplementary tables S3-S4**). Tiling and NFR-focused CRISPRi screens in the GCB-DLBCL cell line Karpas-422 reproducibly identified essential enhancers that in contrast to the MCL cell lines were located exclusively in the 3’ region of the *MYC* locus (**Figure 1B, Supplementary tables S3-S4**). None of our tiling screens showed evidence for essential enhancers in the “Blood Enhancer Cluster” (BENC) that lies at the far 3’ end of the *MYC* TAD and is essential for hematopoietic stem cells, myeloid leukemia, B-cell progenitors, and some B-lymphoblastic leukemias^24–26^, but lacks acetylation in mature B cell populations and lymphomas, indicating that this region is dispensable for mature B-cell lymphomas (**Supplementary Figure S1A**).

We performed CRISPRi screens with the NFR-focused library in a total of two MCL cell lines (one with *MYC* rearrangement) and six GCB-DLBCL cell lines (four with *MYC* rearrangement). Essential gene promoter-targeting sgRNAs showed significant depletion in all eight screens (**Supplementary Figure 1B**). We used the MAGeCK RRA algorithm to score the significance and log2-fold-change of all NFRs across all eight screens (**Figure 1C, Supplementary table S2**), revealing that the dependency landscape of each cell line was dominated by NFRs from specific genomic regions. In all five *MYC*-rearranged cell lines, the most essential NFRs were within the corresponding *MYC* rearrangement partner loci, while MCL and DLBCL cell lines lacking MYC rearrangements were most dependent on the previously mentioned 5’ enhancers and previously uncharacterized 3’ MYC locus enhancers respectively.

### Intra-chromosomal *MYC* rearrangements generate selective *de novo* enhancer dependencies in “hijacked” partner loci

Two cell lines, the MCL cell line JeKo-1 and the GCB-DLBCL cell line OciLy-1, showed intrachromosomal rearrangements between the *MYC* locus and regions of chromosome 8 outside of the *MYC* topologically-associated domain (TAD). In Jeko-1, a chromosomal fusion on the 3’ side of the *MYC* gene linked the *MYC* locus (8q24.21) to 8p23.1, while input chromatin (low-coverage whole-genome sequencing) indicated a copy number gain of the fused regions (**Figure 2A**). CRISPRi revealed strong dependency on a single enhancer NFR within this partner region, NFR 108, that showed no fitness effect in any other cell line. Although NFR 108 is strongly acetylated in JeKo-1, other strongly acetylated elements in the partner locus such as the adjacent NFR 107 showed minimal fitness effect, highlighting the ability of CRISPRi screening to detect selective functional relationships that would not have been predicted by chromatin data alone.

**Figure 2.**
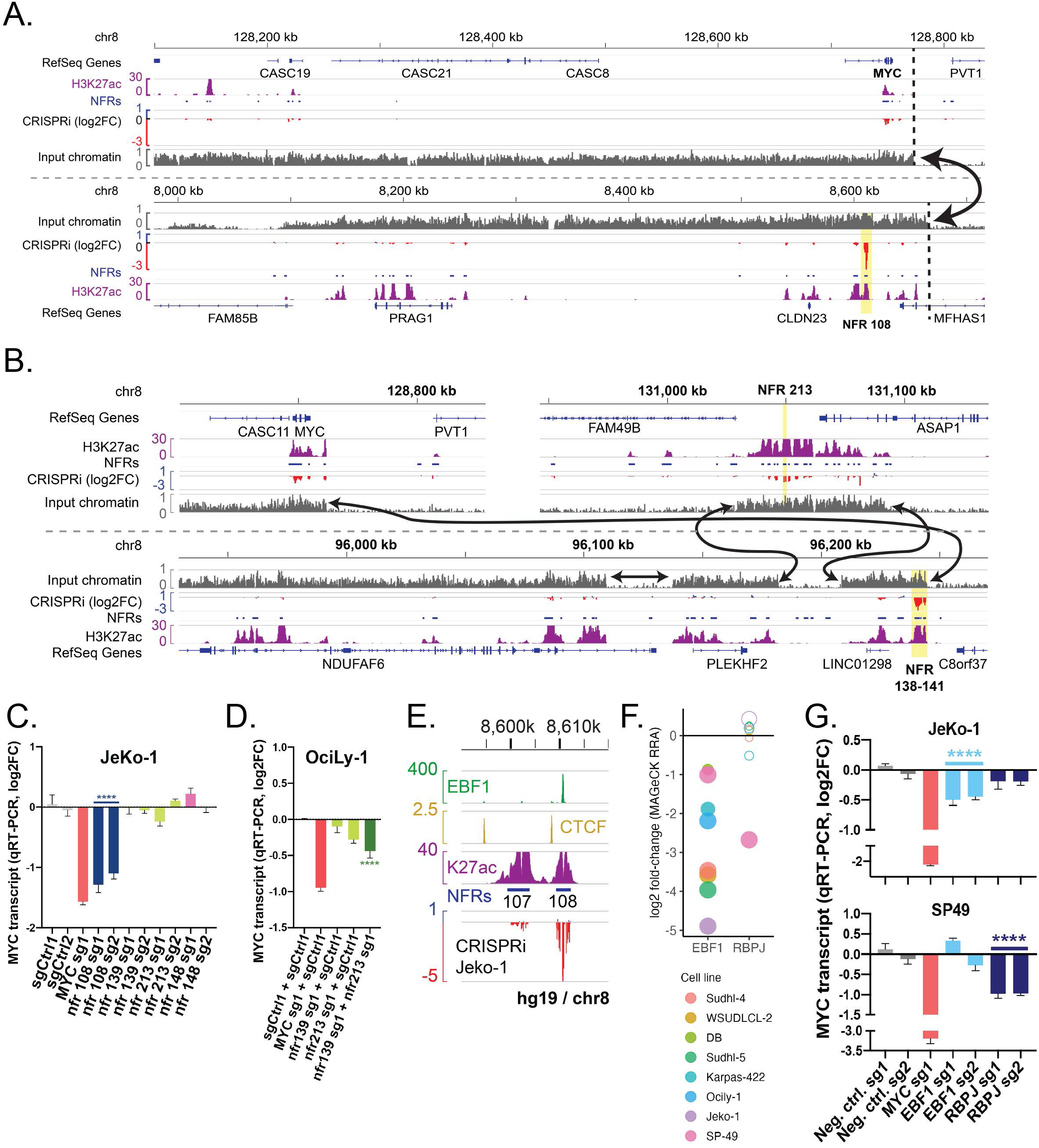
*MYC* activation via intra-chromosomal enhancer-hijacking rearrangements. **A.** H3K27ac ChIP-Seq and CRISPRi screening results for the MYC locus and fusion partner locus in JeKo-1 cells. Input chromatin coverage shows regions of genomic amplification, and arrows connect identified chromosomal fusion breakpoints. **B.** H3K27ac ChIP-Seq and CRISPRi screening results for the MYC locus and fusion partner locus in OciLy-1 cells. Input chromatin coverage shows regions of genomic amplification, and arrows connect identified chromosomal fusion breakpoints. **C.** *MYC* transcript levels (qRT-PCR) after CRISPRi repression of selected enhancers in JeKo-1 cells (**** p<0.0001, t-test of pooled replicates for both nfr 108 sgRNAs versus both control sgRNAs). **D.** *MYC* transcript levels (qRT-PCR) after CRISPRi repression of indicated targets in OciLy-1 cells. Each cell population was transduced with one tagBFP- and mCherry-reporter expressing sgRNA vector. FACS was used to obtain uniform dual-transduced populations prior to doxycycline induction of dCas9-KRAB (**** p<0.0001, t-test of replicates for dual control versus dual enhancer-targeting samples). **E.** Detail of sgRNA CRISPRi scores in JeKo-1 (H3K27ac) and B-lymphoblastoid cell line (CTCF and EBF1) ChIP-Seq data at NFR 108. **F.** Depletion of EBF1 and RBPJ promoter-targeting sgRNAs in 8 lymphoma CRISPRi screens. Circles are sized proportionally to - log2(fdr), with neg|fdr used for log2FC < 0 and pos|fdr used for log2FC > 0. Filled circles represent neg|fdr < 0.01. **G.** MYC transcript levels (qRT-PCR) after CRISPRi knockdown of indicated target genes in selected cell lines (**** p<0.0001, t-test of pooled replicates for both EBF1 or RBPJ promoter-targeting sgRNAs versus both control sgRNAs).

The GCB-DLBCL cell line OciLy-1 bears a more complex *MYC* rearrangement involving two regions of chromosome 8q outside the *MYC* TAD, and with evidence of subsequent chromosomal amplification (**Figure 2B**). Together, these fused regions produce a chain of strongly acetylated enhancers stretching 300 kb from the MYC fusion point. CRISPRi identified one tightly packed cluster of four NFRs located within the space of 5 kb (NFRs 138-141) in rearrangement partner locus 1 (RP1-OciLy-1), as well as weaker and more diffuse essentiality effects across several elements in rearrangement partner locus 2 (RP2-OciLy-1).

To confirm that essential rearrangement partner enhancers activate *MYC* expression, we performed additional CRISPRi studies. Epigenetic repression of NFR 108 lead to a sharp decrease in *MYC* transcript levels after 3 days of dCas9-KRAB induction in JeKo-1, but not OciLy-1 (**Figure 2C, Supplementary Figure S2A**), confirming NFR 108 as a driver of *MYC* expression via enhancer hijacking in JeKo-1. However, the equivalent assay in Oci-Ly1 with single sgRNAs targeting either of the two most essential NFRs identified in the OciLy-1 screen (NFR 139 and NFR 213) failed to significantly repress *MYC* levels, suggesting that these multi-modular enhancers show functional compensation for one another^27, 28^ in the short-term validation assay. Indeed, we found that flow-sorted OciLy-1 populations expressing sgRNAs targeted to NFRs in each of the two partner regions gave substantially stronger *MYC* repression upon dCas9-KRAB induction than repression of either enhancer alone (**Figure 2D**).

Since oncogenic *MYC* activation in JeKo-1 appeared to be selectively driven by a single enhancer (NFR 108) that lacks an essential role in other cell lines, we wondered if this cell line might show increased dependency on trans-regulators of that enhancer. ENCODE ChIP-Seq data from the mature B cell line GM12878 showed that EBF1 was among the factors binding most strongly to NFR 108, which contains a centrally-located EBF1 binding motif (**Figure 2E** **and Supplementary Figure S2B**). While EBF1 is a key component of the mature B cell gene regulatory network and EBF1-targeting sgRNAs were depleted in all cell lines, they showed the strongest depletion in JeKo-1 (**Figure 2F**). The other screened MCL cell line, SP-49, showed unique dependency on *RBPJ*, which encodes the core DNA binding component of the Notch transcriptional complex, consistent with dependency of that cell line on the 5’ B-NDME in the native *MYC* locus. Indeed, knockdown of *EBF1* lead to significant downregulation of MYC transcript levels in Jeko-1 but not SP-49, while *RBPJ* knockdown showed the opposite pattern (**Figure 2G**). These findings suggest that *MYC* rearrangements to specific partner enhancers may shape global *trans*-factor dependencies.

### A recurrent rearrangement places *MYC* under the control of *BCL6* gene-activating mechanisms

The t(3;8)(q27;q24) rearrangement that fuses the *MYC* locus to that of *BCL6*, another key lymphoma oncogene, is a recurrent aberration in GCB-DLBCL (Ohno et al., 2017). A well-characterized distal super-enhancer cluster (*BCL6*-SE) lies 150 kbp upstream of the *BCL6* gene^11, 29–31^. Prior topological evidence indicated that this rearrangement activates *MYC* via enhancer hijacking of *BCL6*-SE elements^11, 21^, but the specific elements responsible for *MYC* activation have not been identified. We performed CRISPRi screens on three lymphoma cell lines, WSU-DLCL2, DB, and SUDHL-4, that bear structurally distinct variants of the *MYC*::*BCL6*-SE rearrangement. We used previously identified genomic breakpoints^20^, 4C-Seq, and optical mapping data to generate parsimonious models for the structure of these rearrangements (**Figure 3A** **and Supplementary Figure S3A-B**), which vary from a simple fusion in WSU-DLCL2 to a complex event including multiple copies of the *BCL6*-SE and additional segments of chromosomes 11 and 1 in SUDHL-4. Topological interactions between the MYC promoter and BCL6-SE were identified in all three cell lines (**Supplementary Figure S3C** and previously shown for WSU-DLCL2^21^). All rearrangement partner regions were included in our NFR-targeting CRISPRi library, thus providing an opportunity to evaluate how complexity affects the function of a recurrent enhancer-hijacking rearrangement.

**Figure 3.**
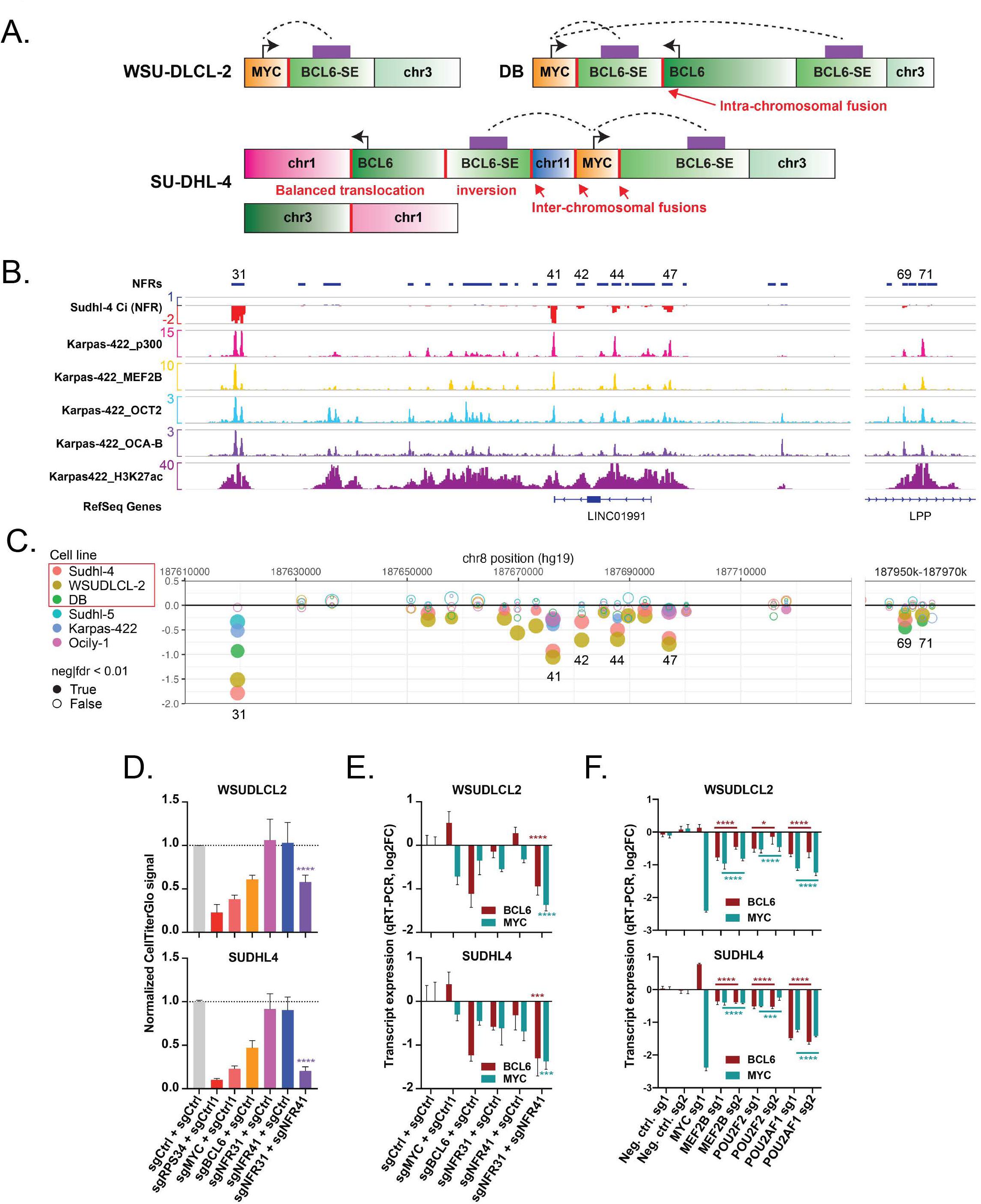
Activation of *MYC* by ternary complex-bound modules of the BCL6 super-enhancer. **A.** Parsimonious models for structural rearrangements involving the *MYC* and *BCL6* locus in three DLBCL cell lines, based on optical mapping and 4C-seq data. Red lines indicate intra- or inter-chromosomal fusions. See Supplementary Figure 3 for details. **B.** CRISPRi sgRNA depletion or enrichment (log2-fold-change) for *BCL6* distal super-enhancer regions in the *MYC::BCL6*-SE+ cell line Sudhl-4. Also shown is ChIP-Seq signal for p300, H3K27ac and ternary complex transcription factors MEF2B, OCT2, and OCA-B in Karpas-422. **C.** Plot of NFR-level sgRNA depletion or enrichment (MAGeCK RRA log2-fold-change, y-axis) for targeted intervals in the BCL6 distal super-enhancer in six GCB-DLBCL cell lines. Circles are sized proportionally to -log2(fdr), with neg|fdr used for log2FC < 0 and pos|fdr used for log2FC > 0. Filled circles represent neg|fdr < 0.01. Note significant and strong fitness signal for ternary factor-bound enhancers in all three *MYC::BCL6*-SE+ cell lines (names in red box). **D.** Chance in viable cellularity for doxycycline-inducible dCas9-KRAB expressing *MYC::BCL6*-SE+ DLBCL cell lines expressing the two indicated sgRNA lentivectors. Cells were transduced with both vectors, sorted for dual tagBFP and mCherry expressing cells, and then grown with or without doxycycline for seven days. Y axis shows the ratio of cellTiterGlo signal for dox treated / untreated cells (**** p<0.0001, t-test of replicates for dual enhancer-targeting sgRNA-transduced versus dual control sgRNA-transduced). **E.** Differential expression of *MYC* and *BCL6* (qRT-PCR) for populations of *MYC::BCL6*-SE+ DLBCL cell lines expressing doxycycline-inducible dCas9-KRAB and two sgRNA vectors as in D (*** p<0.001, t-test of replicates for dual enhancer-targeting sgRNA-transduced versus dual control sgRNA-transduced, color-coded by measured transcript per legend). **F.** Change in MYC expression (qRT-PCR) in *MYC::BCL6*-SE+ DLBCL cell lines expressing doxycycline-inducible dCas9-KRAB after transduction with indicated sgRNAs targeting *MEF2B, POU2F2,* and *POU2AF1* (**** p<0.0001, *** p<0.001, * p<0.05, t-test of pooled replicates for both TF gene promoter-targeting sgRNAs versus both control sgRNAs, color-coded by measured transcript per legend).

CRISPRi screening results in all three *MYC*::*BCL6*-SE+ cell lines were remarkably similar (**Figure 1C** and **Figure 3B-C**). The most essential enhancer in all three cell lines was NFR 31 in the *BCL6*-SE, with other *BCL6*-SE enhancers also showing significant effects. No strongly essential enhancer was identified in the *MYC* locus in these cell lines, nor in either of the additional partner loci in SUDHL-4 (**Figure 1C** **and Supplementary Figure 3D)**. Essentiality effects in all three cell lines were dominated by NFRs that showed selective strong binding of factors previously implicated in enhancer-dependent *BCL6* activation^11, 30^, including the ternary complex members MEF2B, OCT-2, and OCA-B, and the acetyltransferase p300 (**Figure 3B-C**). *BCL6* is also an essential gene in many GCB-DLBCL cell lines, as confirmed by depletion of sgRNAs across the BCL6 promoter region (**Supplementary Figure S3E**), but the fitness effects of sgRNAs targeting the most critical *BCL6*-SE NFRs were stronger in all three *MYC*::*BCL6*-SE+ GCB-DLBCL cell lines than in the other three GCB-DLBCL cell lines. The heightened dependency on *BCL6*-SE elements in *MYC*::*BCL6*-SE+ DLBCL was unlikely to be due to differences in TF binding, as ChIP-Seq binding patterns were similar in *MYC*::*BCL6*-SE+ DLBCL and DLBCL without *MYC* locus rearrangement (**Supplementary Figure S3F**). Assays with single sgRNAs targeting essential *BCL6*-SE NFRs showed only modest effects on *MYC* expression in *MYC*::*BCL6*-SE+ GCB-DLBCL cell lines (**Supplementary Figure S3G**), again suggesting short-term compensatory effects of the multi-modular enhancer, but the combination of sgRNAs targeting two different *BCL6*-SE NFRs resulted in significantly decreased cell growth (**Figure 3E**) and decreased expression of both *MYC* and *BCL6* (**Figure 3F**) in both WSU-DLCL2 and SUDHL-4. CRISPRi knockdown of *MEF2B*, *POU2F2* (OCT2), and *POU2AF1* (OCA-B) encoding the *BCL6*-SE-activating factors^30^, led to similar decreases in both *MYC* and *BCL6* expression in *MYC*::*BCL6*-SE+ cell lines (**Figure 3G**), confirming that the t(3;8)(q27;q24) rearrangement and variants result in lymphomas for which *MYC* expression and ongoing cell growth are highly dependent on the distal cis- and trans-regulatory mechanisms that normally regulate germinal center-specific expression of *BCL6*.

### A novel 3’ enhancer is required for growth and *MYC* expression in GCB-DLBCL lacking *MYC* rearrangement

NFR-targeted CRISPRi screens in non-*MYC*-rearranged GCB-DLBCL cell lines Karpas-422 and Sudhl-5 both identified NFR-195, a site 437 kb downstream (3’) of the *MYC* promoter, as the most essential distal element, with sgRNAs targeting the adjacent NFR 196 showing lesser effects (**Figure 4A-B**). Two other 3’ enhancers, NFR 181 and NFR 202, showed weaker essentiality signal in Karpas-422 tiling and NFR CRISPRi screens, with all three essential enhancers corresponding to regions of locally increased topological interactions with the *MYC* promoter by 4C-Seq (**Supplementary Figure S4A**). NFR 195 was strongly acetylated in two other non-*MYC*-rearranged GCB-DLBCL cell lines, HT and BJAB, and in primary patient GCB-DLBCL biopsies but almost entirely lacked acetylation in *MYC*-rearranged DLBCL cell lines and patient biopsies (**Figure 4A-B** **and Supplementary figure S4B-C**), analogous to the lack of acetylation seen at the 5’ enhancers in *MYC*-rearranged MCL^19^. Low-throughput CRISPRi assays confirmed that NFR 195 is essential in HT and BJAB (**Figure 4C**), and that repression of NFR 195 led to lower *MYC* transcript levels in all four NFR 195-dependent cell lines (**Figure 4D**).

**Figure 4.**
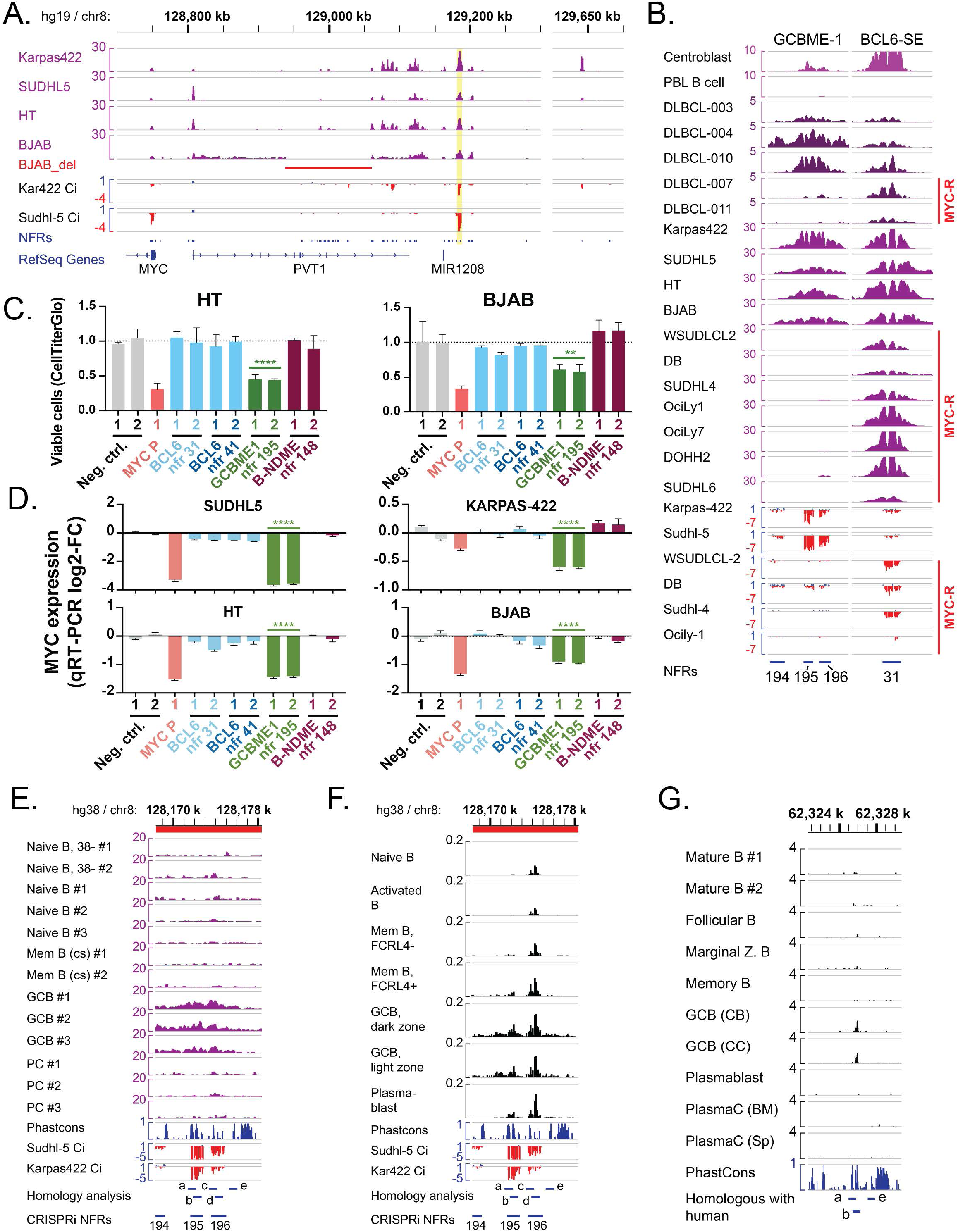
*MYC*-intact GCB-DLBCL is dependent on a 3’ developmental *MYC* enhancer. **A.** H3K27ac ChIP-Seq signal and CRISPRi sgRNA depletion or enrichment (log2-fold-change) from *MYC*-intact GCB-DLBCL cell lines. Yellow highlight shows the location of GCBME-1. **B.** Comparison of H3K27ac ChIP-seq data and CRISPRi sgRNA depletion in GCB-DLBCL cell lines for genomic regions containing GCBME-1 (NFRs 195-196) and the *BCL6*-SE element NFR 31. Also shown at top are H3K27ac ChIP-Seq data from normal human centroblasts and naïve B cells (Farh et al 2014) and GCB-DLBCL patient biopsies (DLBCL-003 to -011; Ryan et al 2015). Note lack of GCBME-1 acetylation in DLBCL biopsies and cell lines with *MYC* rearrangement (MYC-R, indicated at right). **C.** Change in viable cellularity for two additional *MYC*-intact DLBCL cell lines (doxycycline-inducible dCas9-KRAB expressing) after transduction with indicated sgRNAs targeting *MYC* or *BCL6* locus NFRs (** p<0.01, *** p<0.001, t-test of pooled replicates for both NFR 195-targeting sgRNAs versus both control sgRNAs). **D.** Change in *MYC* transcript levels (qRT-PCR) for *MYC*-intact DLBCL cell lines (doxycycline-inducible dCas9-KRAB expressing) after transduction with indicated sgRNAs targeting *MYC* or *BCL6* locus NFRs (**** p<0.0001, t-test of pooled replicates for both NFR 195-targeting sgRNAs versus both control sgRNAs). **E.** H3K27ac ChIP-Seq signal from primary B cell populations (Blueprint consortium) at GCBME-1, showing acetylation specific to germinal center B cells (GCB) at the GCME1 enhancer (NFRs 195-195). Conservation (Phastcons) and CRISPRi signal shown for comparison. “Homology analysis” shows position of intervals evaluated for conservation in other mammalian genomes. **F.** Single-cell ATAC-Seq-derived pseudobulk chromatin accessibility signal for normal tonsil B cell populations (King et al 2021) showing increased chromatin accessibility for germinal center B cells (GCB) at the GCBME-1. **G.** ATAC-seq data from sorted murine B cell populations, centered on sequences syntenic to human NFR 195. Note accessibility specific to germinal center B cell centroblasts (GCB-CB) and centrocytes (GCB-CC). Intervals with substantial homology between hg38 and mm10 are indicated at bottom. Note that sub-regions of human NFR 195 (“a” and “b”) are conserved in mouse, while human NFR 196 (“c” and “d”) lacks an identifiable homolog in mouse.

H3K27ac ChIP-Seq data from normal human lymphoid populations^32^ and cell-type-aggregated single-cell ATAC-Seq data from normal human tonsils^33^ showed selective chromatin acetylation / accessibility at NFR 195/196 in normal germinal center B cells (**Figure 4B, 4E-F and Supplementary Figures S4B-D**). Sequence elements within these NFRs show strong evolutionary conservation, including in mice, and the homologous mouse element also shows GC B cell-specific chromatin accessibility (Immgen project^34^) (**Figure 4G** **and Supplementary Figure 4E**). Together, these findings indicate that NFR 195 / 196 is a physiological germinal center B cell-specific *MYC* enhancer, hereafter ‘germinal center B cell *MYC* enhancer 1’ (GCBME-1), whose function is required to sustain many non-*MYC*-rearranged GCB-DLBCL.

### The ternary complex of MEF2B, OCT2, and OCA-B sustains GCBME-1-dependent *MYC* transcription

*MYC* locus amplification without rearrangement was previously noted by whole-genome sequencing in three DHSig+ primary DLBCL biopsies, one of which (Patient 9) showed further focal amplification of the genomic region containing the GCBME1^9^. We saw no evidence of *MYC* or GCBME-1 amplification in input chromatin from the GCME1-dependent cell lines, although the BJAB cell line shows an interstitial 122 kb deletion within the *PVT1* gene body (**Supplementary figure 4G**). However, *MYC*-intact cell lines such as Sudhl-5 and HT show strong expression of the *MYC* target gene signature comparable to aggressive DHITsig+ primary tumors^4^ and *MYC* transcript levels seen in GCBME-1-dependent cell lines overlap the levels seen in *MYC*-rearranged DLBCL and *MYC*-rearranged Burkitt lymphoma cell lines (**Supplementary figure 4F**), indicating that the GCBME1-dependent cell lines also show hyperactivity of *MYC* transcriptional activation. We examined H3K27ac ChIP-Seq and ATAC-Seq reads mapping to NFR-195 and NFR-196 in the four GCBME1-dependent cell lines, and most nucleotide variants were present at allelic ratios near 50% (**Supplementary Figure 4H**), indicating that enhancer acetylation was biallelic and therefore activated in *trans*.

To look for signatures of differential *trans*-factor activity in GCB-DLBCL cell lines, we expanded our prior genome-wide analysis of B cell cancer enhancer acetylation^10^ to 40 B cell cancer cell lines that spanned the developmental spectrum, including 13 GCB-DLBCL cell lines. Principal component analysis of enhancer acetylation separated cell lines by major biological subtype (**Supplementary figure S5A**). HOMER motif analysis of the enhancer clusters (**Supplementary figure S5B-C and Supplementary Table S5**) confirmed enrichment of motifs for B cell stage-specific TFs in enhancers active in the corresponding tumor subtype. Specific enhancer clusters also revealed motif signatures of genetically activated TFs, such as RBPJ in Notch-rearranged mantle cell lymphoma. Among clusters with specifically increased acetylation in GCB-DLBCL cell lines, we noted that enhancers with strongest activation in the GCBME-1-dependent and *MYC::BCL6*-SE+ cell lines were highly enriched for the OCT2 motif (**Figure 5A**), while enhancer clusters most strongly acetylated in other *MYC*-rearranged GCB-DLBCL showed strong enrichment for the motif of the bHLH factor E2A, but not OCT2. Accordingly, western blot showed high protein levels for the OCT2 ternary complex member MEF2B in GCB-ME1-dependent and *MYC::BCL6*-SE+ cell lines compared to other MYC-rearranged cell lines (**Figure 5B**), while E2A showed generally the opposite pattern (**Supplementary figure 5D**).

**Figure 5.**
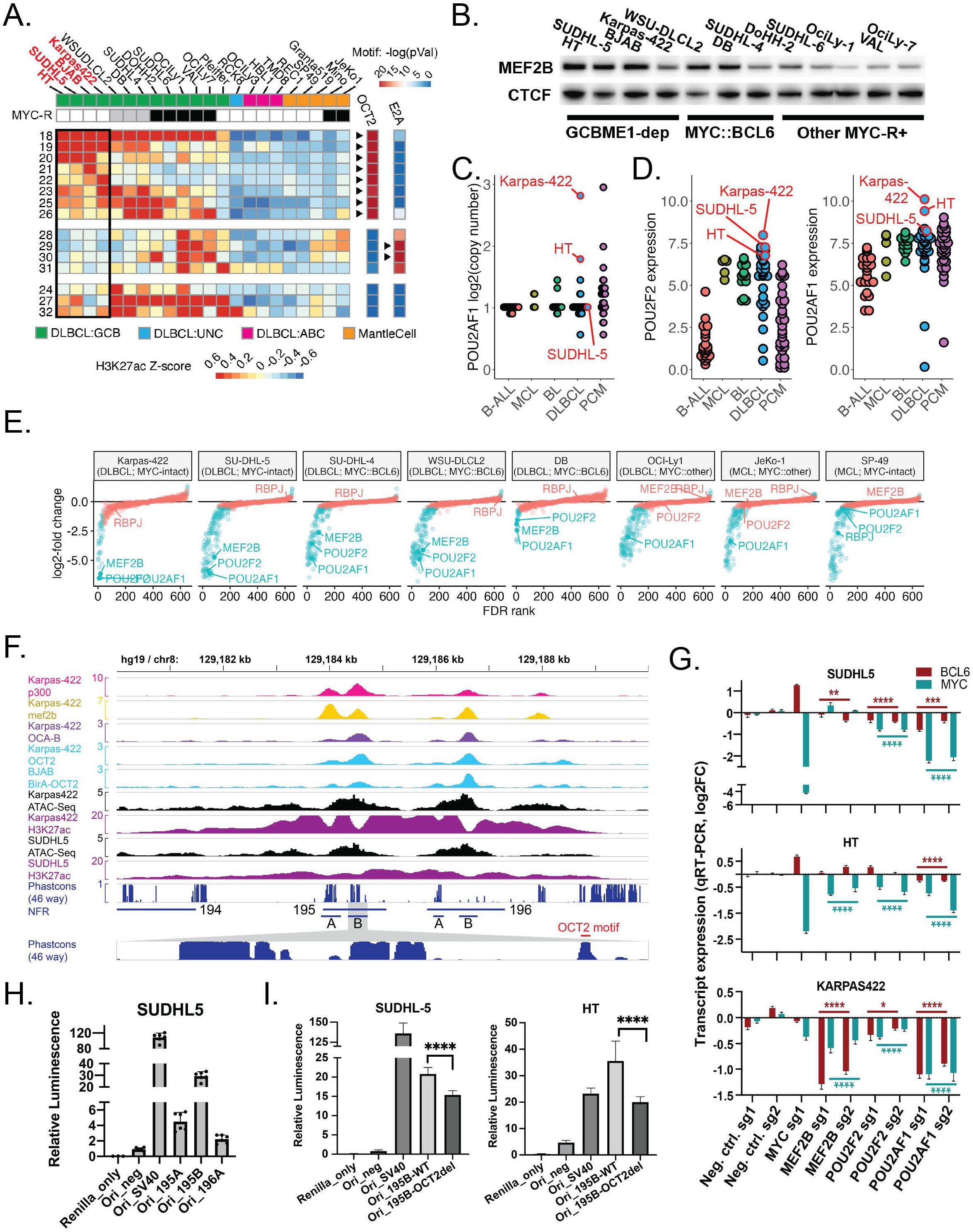
Hyperactivity of the OCT2 / OCA-B / MEF2B complex in GCBME-1-dependent lymphomas. **A.** Heatmap showing differential acetylation of enhancer clusters with increased acetylation in GCB-DLBCL cell lines. *MYC* rearrangement status is coded as follows: white, no rearrangement; gray, positive for MYC::BCL6-SE rearrangement; black, positive for other MYC rearrangement. Heatmaps strips at right show significance of motif enrichment for the OCT2 and E2A (*TCF3*) motifs in HOMER known motif analysis; black arrowheads indicate that a similar motif was identified as the most significantly enriched motif in that cluster by *de novo* motif analysis. See Supplementary Figures 5B-C for the corresponding data for all enhancer clusters and all cell lines. **B.** Western blot of nuclear extracts from GCB-DLBCL cell lines for MEF2B and CTCF (loading control). **C.** Genomic copy gains (log scale) for POU2AF1 in B-cell lymphoma cell lines (DepMap). GCBME-1-dependent cell lines are indicated. **D.** *POU2F2* and *POU2AF1* transcript levels in B-cell lymphoma cell lines (DepMap). GCBME-1-dependent cell lines are indicated. **E.** Plot of sgRNA depletion or enrichment (MAGeCK RRA log2-fold-change) for optimized gene promoter-targeting sgRNAs in lymphoma CRISPRi screens (B cell expressed transcription factors, B-cell-specific essential genes, and pan-essential genes). Target intervals are ranked along the X axis by lowest negative selection FDR p-value (left to right, intervals with log2-fold change <0) or lowest positive selection FDR p-value (right to left, intervals with log2-fold change >0). Genes with abs(log2FC) > 0.5 and fdr < 0.05 are colored blue. TF genes of interest are indicated. **F.** Detail of ATAC-Seq, H3K27 ChIP-Seq, and ChIP-Seq for the OCT2, OCA-B, MEF2B, and p300 in the GCBME1 enhancer cluster. CRISPRi screen NFR intervals are as indicated, with conserved subregions A and B of NFRs 195 and 196 indicated below. Inset at bottom shows further detail of the Phastcons conserved element track for sub-region 195B (see Supplementary Figure 5G for further detail), with the position of the conserved OCT2 motif indicated. **G.** Change in MYC expression (qRT-PCR) in GCBME1-dependent DLBCL cell lines expressing doxycycline-inducible dCas9-KRAB after transduction with indicated sgRNAs targeting *MEF2B, POU2F2,* and *POU2AF1* (**** p<0.0001, *** p<0.001, ** p<0.01, * p<0.01, t-test of pooled replicates for both TF gene promoter-targeting sgRNAs versus both control sgRNAs, color-coded by measured transcript per legend). **H.** Transcriptional reporter assay in SUDHL-5 cells for sub-regions of the GCBME-1 cloned into the STARR-Seq_Ori luciferase vector. Firefly luciferase signal for each sample was normalized to internal transfection control (Renilla luciferase signal) and then to the normalized signal for the no-insert control vector (Ori_neg). **I.** Transcriptional reporter assay in the indicated GCBME-1-dependent cell lines for constructs bearing the NFR 195B enhancer region, with or without OCT2 motif deletion, normalized as in H.

The genes encoding OCT2 (*POU2F2*, chr19q13.2) and its coactivator OCA-B (*POU2AF1*, chr11q23.1) lie within regions of frequent large-scale copy gain (Chapuy et al., 2018) and less frequent focal amplification (Hodson et al., 2016) in DLBCL. Strikingly, three of the four GCME1-dependent GCB-DLBCL cell lines contained focal copy gains affecting of one of these genes, with *POU2AF1* amplified in Karpas-422 and HT, (**Figure 5C**), and *POU2F2* showing a focal copy gain in BJAB (Hodson et al., 2016). Western blot confirmed higher expression of the corresponding proteins in gene-amplified cell lines (**Supplementary figure 5D**). Overall, the three GCME1-dependent cell lines with RNA-Seq data in DepMap show higher expression of *POU2F2* and *POU2AF1* transcripts than most other DLBCL cell lines (**Figure 5D**). Gene promoter-targeting sgRNAs in our CRISPRi screens confirmed that *POU2F2*, *POU2AF1*, and *MEF2B* were among the most essential TF genes in both GCBME-1-dependent and *MYC::BCL6-SE*+ DLBCL cell lines (**Figure 5E**), while *POU2F2* and *MEF2B* were dispensable for OciLy-1.

Strong ChIP-Seq binding peaks for p300 acetyltransferase and all three members of the OCT2 / OCA-B / MEF2B ternary complex were noted at multiple subunits of the GCBME-1 enhancer cluster (**Figure 5F, Supplementary Figure 5E**), suggesting that high expression of these factors contribute to MYC activation through the native enhancer. Knockdown of *POU2F2*, *POU2AF1*, and *MEF2B* by CRISPRi and shRNA led to decreased *MYC* transcript and protein levels in GCBME-1 dependent cell lines (**Figure 5G**), with *POU2AF1* knockdown showing the strongest and most consistent effects. Interestingly, while effects of ternary complex factor knockdown led to closely correlated decreases in *MYC* and *BCL6* expression in MYC::BCL6-SE+ cell lines (**Figure 3F**), we saw more divergent effects in two of the GCBME-1-dependent cell lines, with stronger effects of POU2AF1 knockdown on *MYC* versus *BCL6* expression (**Figure 5G**).

We next sought to identify conserved elements within GCBME-1 that might mediate direct binding by the ternary complex. We divided NFRs 195 and 196 into sub-regions based on dips of H3K27ac signal, peaks of TF ChIP-Seq signal and clusters of evolutionarily conserved elements (**Figure 5F** and **Supplementary Figure 5G-H**). Of these, 195B corresponded to the region of strongest sgRNA depletion in all Karpas-422 and SUDHL-5 CRISPRi screens, showed strong binding of all three TFs and p300, and unlike regions 196A&B was conserved in nearly all mammals. Luciferase reporter constructs confirmed that region 195B was a strong transcriptional driver in SUDHL-5 cells, while 195A and 196A showed weaker effects (**Figure 5H**). Deletion of an evolutionarily conserved OCT2 binding motif in NFR 195B significantly reduced transcriptional reporter activity in two GCBME-1-dependent cell lines (**Figure 5I**), supporting NFR 195B as a key response element that confers ternary complex-dependent *MYC* activation.

### A recurrently deleted *cis*-element inhibits *MYC* activation by native and hijacked enhancers

Non-*MYC*-rearranged high-grade GCB-DLBCL patient samples are enriched for focal deletions removing the promoter (*PVT1*-P) of the non-coding RNA gene *PVT1* (Hilton et al., 2019). Since *PVT1*-P was previously shown to act in *cis* as a competitive antagonist of *MYC* activation by 3’ enhancers in breast cancer cells (Cho et al., 2018), loss of *PVT1*-P activity could represent a mechanism of increased *MYC* activation by native B cell enhancers. Indeed, both our tiling and NFR-focused CRISPRi screens identified significant *enrichment* of sgRNAs targeting *PVT1*-P in GCBME-1-dependent cell lines (**Figure 6A-B**), with weaker effects seen in MCL cell lines driven by 5’ MYC enhancers. Importantly, *PVT1*-P also appeared to affect regulation of *MYC* by heterologous 3’ enhancers, as we also saw a fitness-enhancing effect of *PVT1*-P silencing in the *MYC::BCL6*-SE+ cell lines SUDHL-4 and WSU-DLCL2, in which the MYC locus breakpoint is downstream of *PVT1*-P (**Figure 6A-B, Supplementary Figure S6A**). In contrast, we saw no growth-promoting effect of *PVT1*-P silencing in OCI-Ly1, Jeko-1, or DB, in which 3’ fusion breakpoints occur between the *MYC* gene and *PVT1*-P, excluding the latter from the fusion chromosome. We confirmed that CRISPRi silencing of the *PVT1* promoter increases *MYC* transcript levels in cell lines dependent on the 3’ GCBME-1 (**Figure 6C**), with most *PVT1*-P-targeting sgRNAs also increasing *MYC* expression in cell lines dependent on 5’-enhancers or on a rearrangement placing the *BCL6*-SE in *cis* with *PVT1*-P and *MYC* (**Figure 6D, Supplementary Figure S6B**).

**Figure 6.**
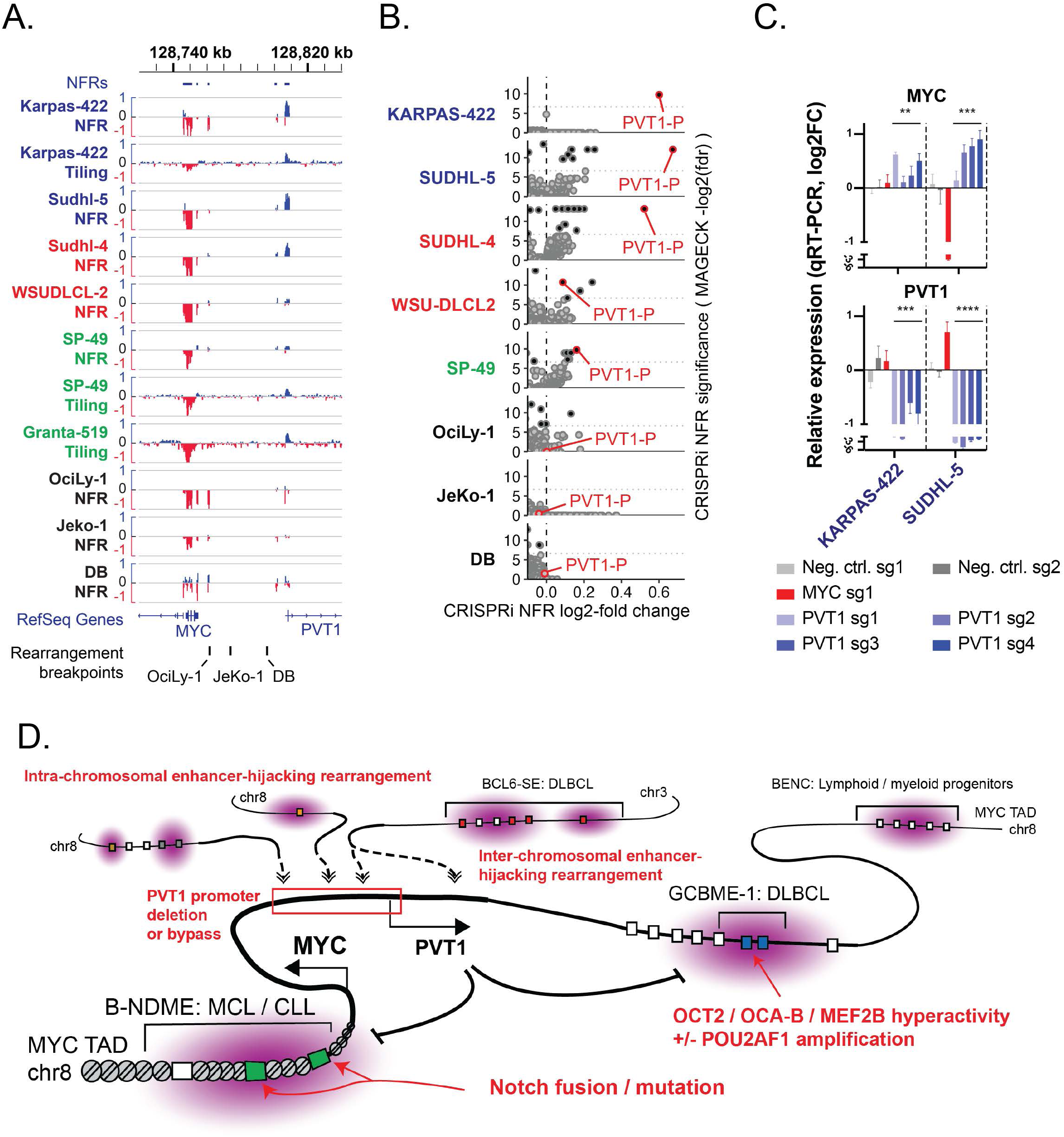
The recurrently deleted *PVT1* promoter is a cis-repressor of GCBME-1 dependent *MYC* activation. **A.** Depletion or enrichment (log2-fold-change) for sgRNAs targeting the *MYC* and *PVT1* promoter regions in *MYC* locus tiling (20 sgRNA sliding window) and NFR-focused CRISPRi screens (log2-fold-change of individual sgRNAs). Cell line names are color-coded as follows: Blue, *MYC*-intact GCB-DLBCL; Red, *MYC::BCL6*-SE+ GCB-DLBCL with the breakpoint downstream of the *PVT1* promoter; Green, *MYC*-intact MCL; Black, DLBCL and MCL cell lines with *MYC* rearrangement breakpoint between the *MYC* gene and the *PVT1* promoter. Rearrangement breakpoints shown at bottom are from Ryan et al 2015 and Chong et al 2018. **B.** Volcano plots of MAGeCK analysis on NFR CRISPRi screens (cropped to highlight enriched regions), showing enrichment of sgRNAs targeting the *PVT1* promoter (nfr 156). Cell line names are color-coded by subtype and rearrangement as in A. **C.** *PVT1* and *MYC* transcript levels (qRT-PCR) in GCBME-1-dependent cell lines (doxycycline-inducible dCas9-KRAB expressing) after transduction with indicated sgRNAs targeting the *MYC* or *PVT1* promoters (**** p<0.0001, t-test of pooled replicates for both NFR 195-targeting sgRNAs versus both control sgRNAs) **D.** Summary of *MYC* activation mechanisms identified in *MYC*-rearranged and MYC-intact mature B cell lymphoma cell lines.

## Discussion

*MYC* is a master regulator of gene programs central to rapid cellular proliferation and anabolic metabolism in diverse tissues^35^, but its upstream mechanisms of transcriptional activation are highly lineage- and state-specific, with this regulatory specificity encoded by a massive diversity of tissue-specific enhancers located throughout the 3 Mbp *MYC* topological domain^12, 19, 24, 26, 36, 37^. Precise regulation of *MYC* is critical in normal germinal center B cells, where transient upregulation of *MYC* is critical for driving repeated cycles of immunoglobulin gene mutation, selection, and affinity maturation^38^. Common polymorphisms associated with altered risk for mature B-cell leukemias and lymphomas occur in the vicinity of state-specific B cell *MYC* enhancers^39–41^, and slight *MYC* transcript stabilization conferred by mutant BTG1 leads to profound effects on B cell selection and lymphomagenesis^42^. Although the 3’ region of the *MYC* locus downstream of the *PVT1* promoter has been noted to contain numerous candidate mature B-cell enhancers in mice^43^ and humans^11^ based on chromatin state and topological interactions with the *MYC* promoter, to our knowledge the GCBME-1 enhancer is the first element in this region that has been functionally proven to drive *MYC* expression in B cells. While GCBME-1 is unlikely to be the only distal element that controls the highly dynamic regulation of *MYC* transcription during the normal germinal center reaction, our identification of this evolutionarily conserved element may open new opportunities for functional elucidation of *MYC* transcriptional in germinal center biology and immunity.

Identification of GCBME-1 as a common dependency of GCB-DLBCL cell lines lacking *MYC* rearrangements suggests alternative mechanisms that may drive increased *MYC* expression in *MYC*-intact DLBCL. The *POU2AF1* gene is located on chromosome 11q, and is often included in the recurrent chromosomal gain seen in “high grade B-cell lymphoma with 11q aberration”, a lymphoma subtype defined by a high-grade Burkitt lymphoma-like morphology, phenotype, and gene expression program, but with a characteristic pattern of amplification and deletion on chromosome 11q and no *MYC* locus rearrangement^44^. The GCBME-1-dependent cell lines SUDHL-5 and HT bear characteristic 11q abnormalities, and have been previously characterized as models for this entity^44^, although only HT shows *POU2AF1* amplification. Further work is indicated to determine whether other genes dysregulated by the recurrent 11q aberration may contribute to *MYC* activation by GCBME-1. Although *MYC* protein levels in GCBME-1-dependent lymphoma cell lines tended to be lower than in many *MYC*-rearranged cell lines (**Supplementary figure 5D**), our findings show that GCBME-1 activity and *MYC* transcription are rate-limiting steps for the fitness of these lymphomas, suggesting that targeting mechanisms of GCBME-1 activation could be a useful therapeutic approach.

Our study also provided novel insights into mechanisms of *MYC* activation via rearrangement to non-immunoglobulin loci, confirming that such events can create new cancer-specific dependencies on individual enhancer elements or super-enhancer arrays in partner loci, as well as their cognate trans-factors. The *BCL6* locus appears to be the most common non-immunoglobulin partner locus of *MYC* rearrangements, resulting in “pseudo-double-hit” rearrangements that can be mis-interpreted as separate rearrangements of *MYC* and *BCL6* by standard clinical FISH studies. Our findings confirm that the MEF2B / OCT2 / OCA-B ternary complex-bound subunits of the distal *BCL6*-SE / locus control region are the critical drivers of *MYC* activation in such rearrangements, and that these *MYC::BCL6*-SE+ lymphomas retain strong dependency on the highly germinal center state-specific *BCL6*-SE, while expression of *BCL6* itself can be released from normal regulatory controls by diverse DLBCL-associated lesions, including *BCL6* “promoter swap” rearrangements and mutations in the BCL6 first intron / intragenic super-enhancer^45, 46^. By re-wiring *MYC* regulation to enhancers that rely on distinct combinations of stage-specific trans-factors, specific *MYC* rearrangements may constrain the resulting lymphoma to specific developmental states and epigenetic dependencies.

## Acknowledgements

R.J.H.R was supported by grants from the National Cancer Institute (K08-CA-208013 and R01-CA-245059), the American Society of Hematology (Scholar Award), and The V Foundation for Cancer Research (V2018-001). J.M.E. was supported by NIH NHGRI (K99HG009917 and R00HG009917), the Harvard Society of Fellows, Gordon and Betty Moore, and the BASE Research Initiative at the Lucile Packard Children’s Hospital at Stanford University. The authors wish to thank Yuanyuan Chang and Joyce Lee at Bionano Genomics for assistance with optical mapping data generation and analysis, and Bradley E. Bernstein, Mark Y. Chiang, and Sami N. Malek for helpful feedback on the manuscript.

## Author contributions and competing interests

A.I. and A.G. performed research, analyzed data, and wrote the paper, R.K., A.A., D.R., J.G., C.N.H., J.S.R., M.W., M.R. and R.A. performed research, I.M.G supervised research and contributed vital new reagents, C.P.F. and J.M.E. supervised research and analyzed data, T.S., A.R., and M.P.C. analyzed data, R.J.H.R. designed research, supervised research, analyzed data, and wrote the paper.

J.M.E. is an equity holder and consultant for Martingale Labs, a provider of polygenic genomic risk scores for healthcare that was not directly or indirectly involved in this research.

## Supplementary Methods

### ChIP-Seq

For H3K27ac ChIP-Seq, five million cells were cross-linked in PBS + 1% formaldehyde for 10 minutes with periodic inversion, quenched for 5 minutes with glycine added to a final concentration of 125 mM, washed twice in PBS plus protease inhibitors (PI; Roche cOmplete), and pellets were frozen at -80C. Pellets were thawed and nuclei isolated via a cytoplasmic lysis buffer (20 mM Tris-HCl pH 8.0, 85 mM KCl, 0.5% NP 40 + PI) and centrifugation at 400g for 5 min at 4°C. Nuclei were resuspended in cold SDS lysis buffer (0.3% SDS, 10mM EDTA, 50mM Tris-HCl, pH 8.1 + PI), homogenized by passage through a 27g needle. Sonication was performed with a Q800R2 Sonifier (QSonica) set to amplitude 70 alternating between 45s on and 15s off for a total of 8min 30s on. Samples were then diluted 1:3 in ChIP dilution buffer (0.01% SDS, 1.1% Triton X-100, 1.2mM EDTA, 16.7mM Tris-HCl, pH 8.1, 167mM NaCl +PI), and rotated at 4°C overnight with 2 ug of antibody (H3K27ac, #39133, Active Motif). Chromatin complexes were bound for 4 hours on Protein G Dynabeads (Thermo). Subsequent washing, DNA elution, purification, and Illumina library preparation steps were performed as previously described^1^.

For transcription factor ChIP-Seq, the protocol used was previously detailed^2^ and differs from that above as follows: 20 million cells were crosslinked in media + 1% formaldehyde 37°C, sonication was performed in higher SDS buffer (1% SDS, 10mM EDTA, 50mM Tris-HCl, pH 8.1 + PI) with a Branson probe sonifier for sonication. 2-5 ug of antibody was used per immunoprecipitation.

### ATAC-seq

ATAC-Seq was performed as previously described^1^. Nuclei were isolated from 50,000 cells for each sample using Nuclei EZ prep-Nuclei Isolation Kit (Sigma-Aldrich). The transposition reaction mix (25 μL of 2× TD buffer, 2.5 μL of Tn5 transposase (Illumina), 15 μL of PBS and 7.05 μL of nuclease-free water) was added to nuclei and incubated at 37°C for 1 hour in an orbital shaker at 300 RPM. 50 μL Qiagen buffer PB was added to each sample to stop the reaction and DNA was isolated with AMPure XP beads (Beckman Coulter). Fifteen cycles of PCR were performed with transposed DNA using the dual index primers and NEBNext PCR Master Mix, followed by AMPure XP purification. After quantification and fragment size analysis, libraries were sequenced on Illumina NextSeq.

### NFR CRISPRi library design

To design the NFR-focused sgRNA library, we defined target regions within the MYC locus to include the MYC / PVT1 promoter region (chr8_MYC_genic) as well as 5’ (chr8_MYC_5’enh) and 3’ (chr8_MYC_3’enh) regions that include all strongly acetylated elements in the subject cell lines and interact topologically with the *MYC* promoter (see *MYC* promoter 4C-Seq data from the Karpas-422 cell line in this study and published 4C-Seq data from REC-1 cell line^3^. We covered the entire topologically-associated domain (TAD) upstream of BCL6 (chr3_BCL6_5’enh). We also included two additional rearrangement partner (RP) regions linked to the MYC and BCL6 genes in the SUDHL4 cell line based on optical mapping and 4C-Seq (chr11_RP1_SUDHL4 and chr1_RP2_SUDHL4), and regions of chr8 rearranged to the MYC locus in the Jeko-1 (chr8_RP_JeKo1) and OciLy1 (chr8_RP1_OCILy1 and chr8_RP2_OCILy1) based on PEAR-ChIP. Within these regions, ATAC-Seq peaks from 26 DLBCL and MCL cell lines were used to define candidate nucleosome-free regions (NFRs), which were extended to 500 bp from the peak center. The estimated enhancer “activity score”^4^ was calculated as the geometric mean of normalized H3K27ac and ATAC-Seq reads in each peak for each cell line and peaks with activity <15 were discarded. Remaining peaks across the 26 cell lines were merged into a consensus set of NFRs with high predicted enhancer activity in at least one cell line; overlapping peaks were merged.

All candidate sgRNAs within the target region NFRs were identified and scored with FlashFry ^5^ and sgRNAs were retained that met the following scoring criteria: Doench2014OnTarget > 0.1, Hsu2013 > 50, JostCRISPRi_specificityscore > 0.1, dangerous_GC == “NONE”, dangerous_polyT == “NONE”, dangerous_in_genome == “IN_GENOME=1”, otCount < 500. Remaining sgRNAs in each interval were downsampled to not more than 35 per 1 kb.

The final libraries also contained 646 sgRNAs covering extended tiling regions over the *MYC* and *BCL6* promoters and proximal gene bodies, and 392 guides targeting non-enhancer control regions in the *MYC* and *BCL6* loci. We also included CRISPRi-optimized sgRNAs (5 per gene) from the Dolcetto library^6^ targeting the promoters of 50 pan-essential genes, 38 genes selectively essential in B-cell lymphoma based on DepMap CRISPR knockout screens^7^, and 575 transcription factor genes expressed in B cell cancer cell lines. 866 non-targeting sgRNAs were included as negative controls.

Pools of oligos with the sequence 5’-TATCTTGTGGAAAGGACGAAACACCG-{20-mer-seed}-GTTTAAGAGCTATGCTGGAAACAGCATAG-3’ were ordered from CustomArray or Twist Biosciences. 24ng of the oligo pool was PCR amplified using NEBNext master mix and 1uM each of sgRNA_Library_Fwd/Rev primers. PCR products were purified with 1.5X AMPureXP beads and cloned into BsmBI-digested sgOpti by Gibson assembly with NEBuilder HiFi DNA Assembly master mix per manufacturer’s protocol and a 5-fold molar excess insert to vector ratio. 1 uL of assembly reaction was electroporated with a Bio-rad Gene Pulser Xcell system into Endura competent cells, and culture outgrowth, determination of transfection efficiency, and plasmid purification were performed as described^8^.

### Optical mapping

Fresh-frozen cell line pellets were sent to Bionano Genomics for high-molecular weight DNA extraction, direct labelling with DLE-1, DNA backbone counterstaining, and imaging on a Saphyr chip. Optical map images were converted to BNX format after which de novo assembly and structural variant detection were performed using Bionano Genomics pipelines. Structural variant read maps involving the MYC, BCL6, and partner loci were visualized with BioNano Access software v1.7.2.

### 4C-Seq

4C-Seq analysis was performed on Karpas-422, DB, and SUDHL-4 cells as previously described^3, 9^, with the 4-cutter restriction enzymes DpnII (primary) and CviQIF (secondary). Viewpoint primers for the *MYC* promoter and the *BCL6*-SE element NFR 31 are listed in Supplementary Table 1. Heatmaps of 4C interaction frequencies across a range of resolutions in genomic loci of interest were generated with 4C Seq Pipe^10^. To visualize interactions across the *MYC::BCL6*-SE fusion in DB and Sudhl-4, 4C Seq pipe was run with a custom DpnII-CviQIF-digested hg19 reference containing a ch8 / chr3 fusion chromosome (chr8 position 1-128,760,000, followed by chr3 position 187,480,000 to terminus).

### Genome-wide enhancer acetylation and motif analysis

Genome-wide clustering of enhancer modules / NFRs based on differential acetylation was performed on ATAC-Seq and H3K27ac ChIP-seq datasets as previously described for 26 cell lines, ^1^, here expanded to 40 B cell cancer cell lines. Briefly, ATAC-Seq peaks from all 40 cell lines were resized to 200 bp, then merged into a union set of consensus 200 bp intervals via GenomicRanges “reduce” and “resize”. Next, read-count normalized H3K27ac ChIP-seq signal was quantified for each cell line in a 1,000 bp window around each peak. Intervals located < 2 kb upstream or < 1 kb downstream of an annotated TSS and intervals associated with low H3K27ac signal in all cell lines were removed, and the matrix of cell line acetylation values for each peak were square root transformed, centered, scaled and used for principal component analysis of cell lines and and k-means clustering of enhancers (k=60). HOMER findMotifsGenome was run on each resulting enhancer cluster for known and de novo motif analysis, with the set of all enhancers used as a background (option -b).

### Enhancer sequence conservation and motif analysis

To systematically identify conserved intervals in mm10 that correspond to hg19 NFRs, we used a UCSC hg19-mm10 genomic alignment net file to identify the mm10 position corresponding to the center-most base in each hg19 NFR (merged ATAC-Seq peaks as described above) that is mappable to mm10. Mapped intervals were then extended to 200 bp for visualization. To extract conserved sequences from multiple vertebrate species corresponding to the GCBME-1 NFRs, evolutionarily conserved sub-regions were manually defined based on clusters of predicted evolutionarily conserved elements in hg38 (Phastcons vertebrate 100-way). We then extracted ungapped sequences aligned to the hg38 intervals consisting of the longest mappable region with base matching ratio > 0.1 for each of 70 vertebrate species for which pairwise MULTIZ chain files to hg38 were available. HOMER known motif analysis was then applied to the syntenic interval for each species, and specific motifs and motif families of interest were selected for visualization.

### Enhancer variant analyses

Sequence variants in the GCME enhancer (hg19-chr8:129,180,000-129,190,000) were identified with bcftools mpileup in DLBCL H3K27ac ChIP-Seq bam files. The ratio of reference to alternate bases was calculated for loci represented in at least 10 reads.

**Supplementary Figure 1.**
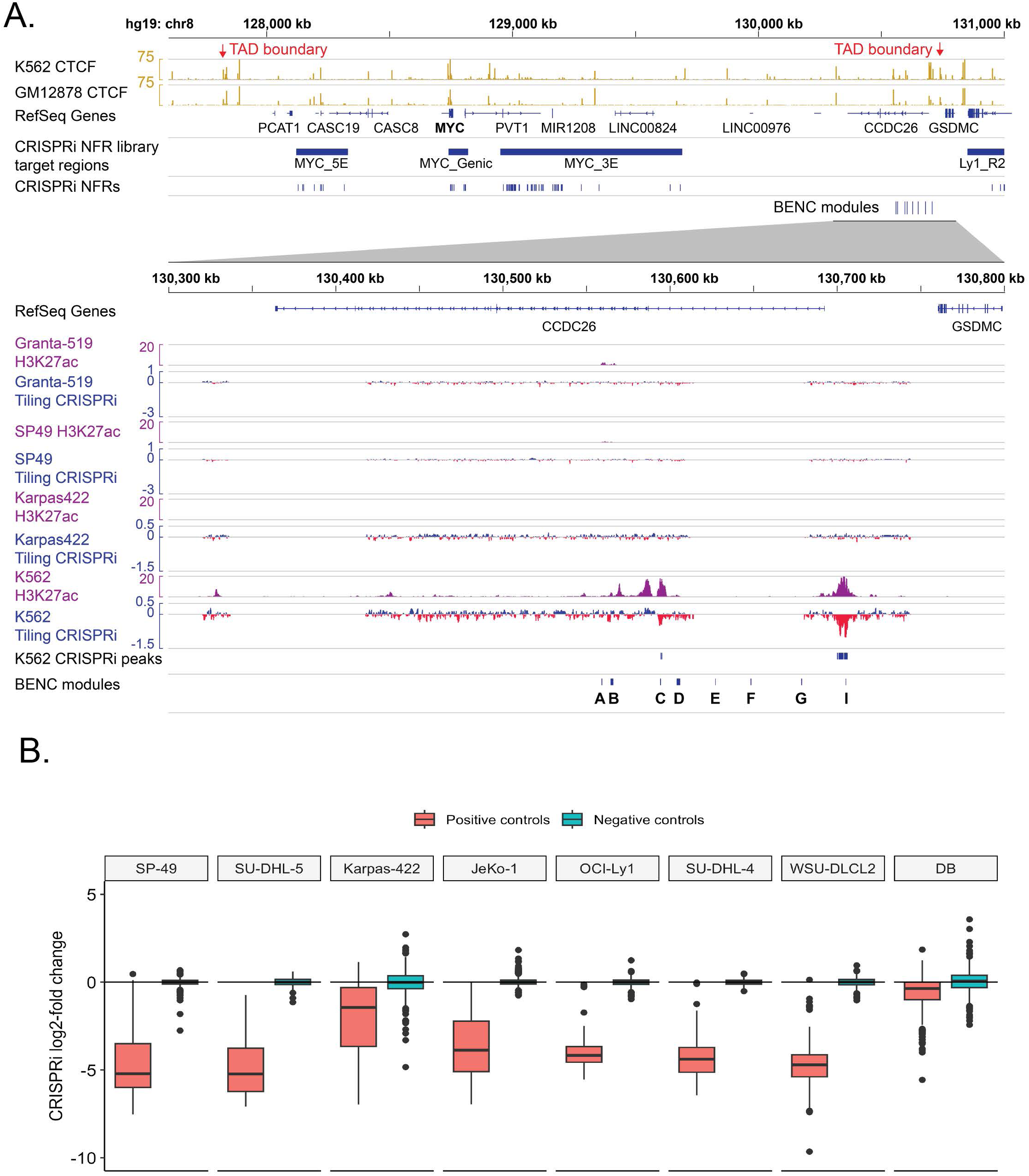
**S1A.** Top: Complete *MYC* locus, showing regions targeted by the NFR CRISPRi library at bottom. The boundaries of the *MYC* topologically associated domain (TAD) correspond to CTCF peaks (ENCODE ChIP-Seq) indicated by arrows, as identified by Hi-C^11^. Note that one of the rearrangement partner regions identified in OciLy-1 (Ly1_R2) lies just outside the *MYC* TAD. Bottom: Detail of H3K27ac ChIP-Seq data and CRISPRi scores (log2-fold change, 20 sgRNA sliding window) from tiling CRISPRi screens performed in three mature B-cell lymphoma cell lines (this study) and K562 cells^11^ in the region of the blood enhancer cluster (BENC) previously shown to play a key role in early hematopoietic and lymphoid development. **S1B.** Summary of CRISPRi scores for positive control (pan-essential gene promoter targeting) and negative control (non-targeting) sgRNAs for each of 8 NFR-targeting CRISPRi screens. Note that essentiality effect sizes in the DB and Karpas-422 screens may be underestimated due to lower CRISPRi efficiency than other cell lines.

**Supplementary Figure 2.**
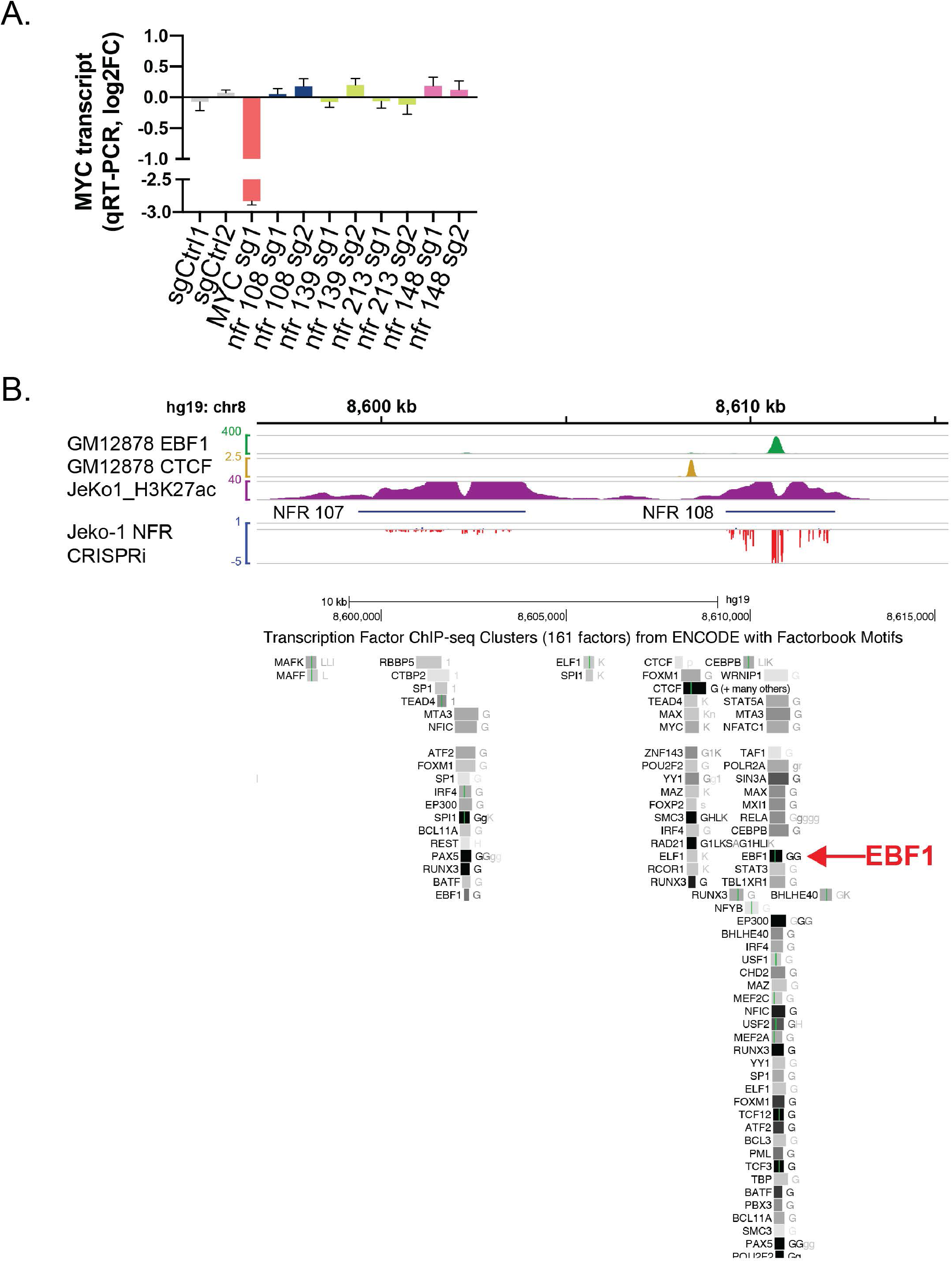
**S2A.** *MYC* transcript levels (qRT-PCR) after transduction with sgRNAs targeting selected NFRs in OciLy-1 cells with doxycycline-inducible KRAB-dCas9. **S2B.** ChIP-Seq signal and CRISPRi sgRNA depletion / enrichment in the indicated cell lines in the vicinity of NFR 108. Selected TF binding peaks are shown at bottom as identified by ENCODE ChIP-Seq. Green lines indicate matching Factorbook motifs for the corresponding factor and grayscale corresponds to peak strength (see UCSC genome browser for schema). Peaks labelled with “G” were identified in EBV-transformed peripheral blood B cells (GM12878).

**Supplementary Figure 3.**
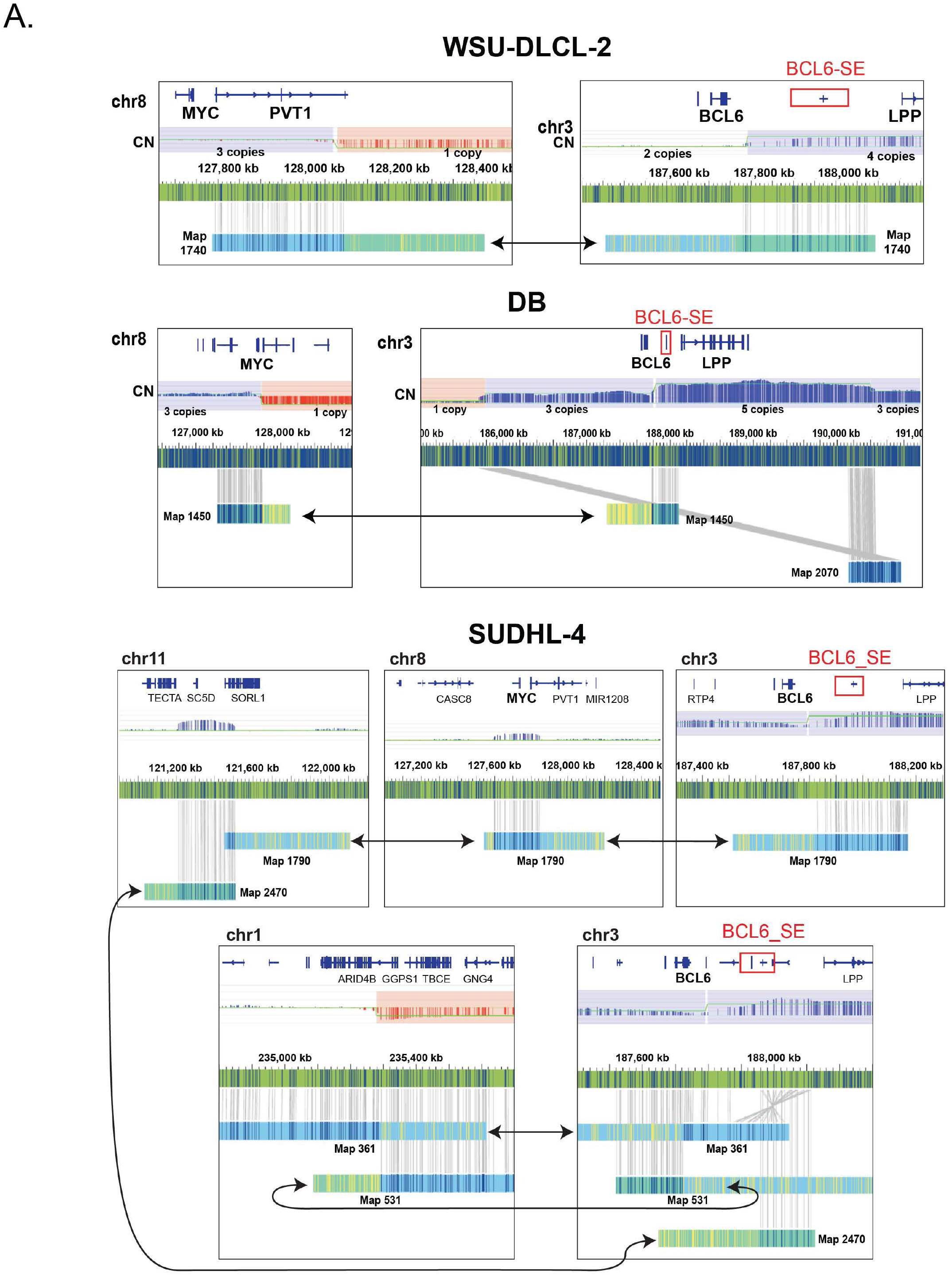
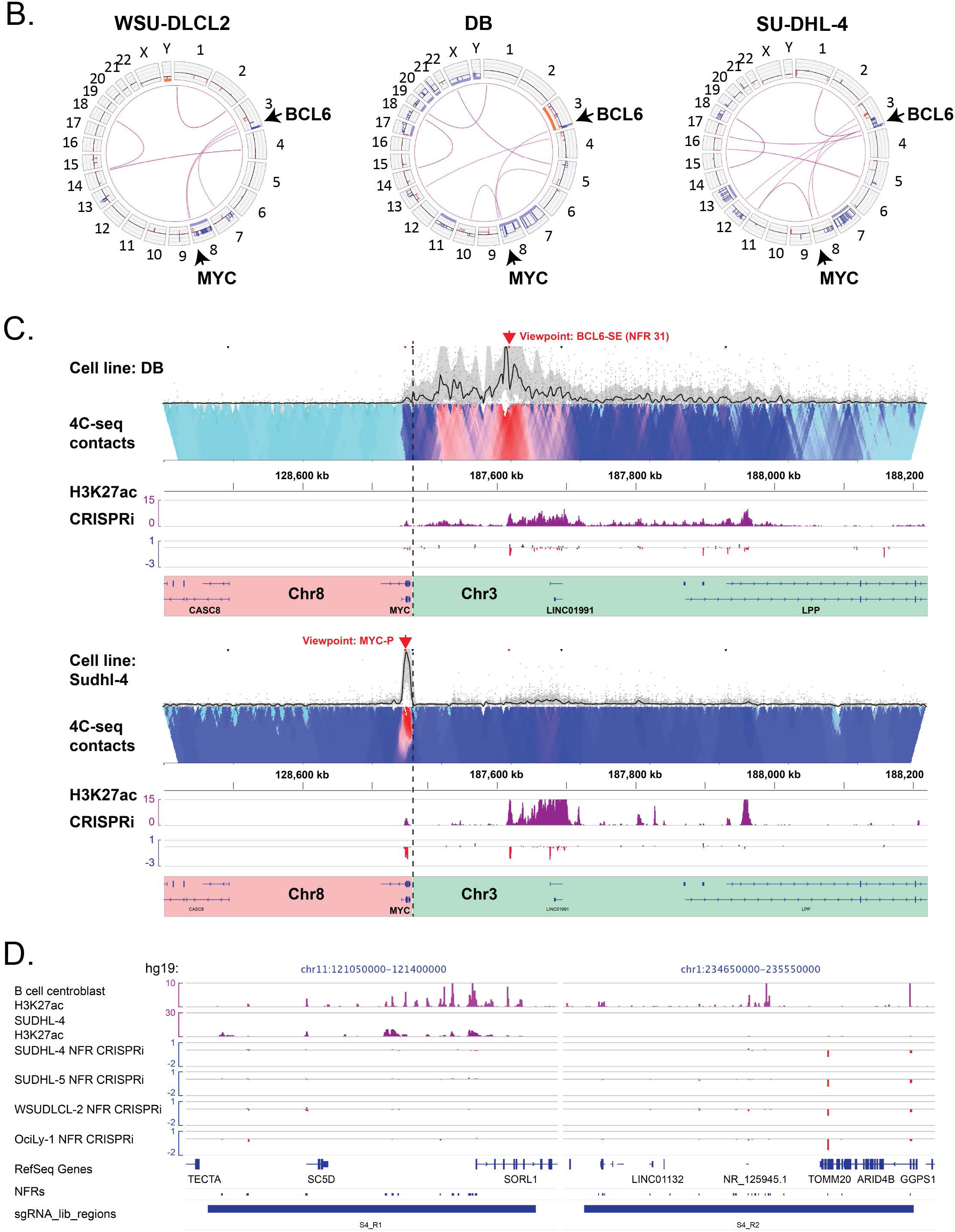
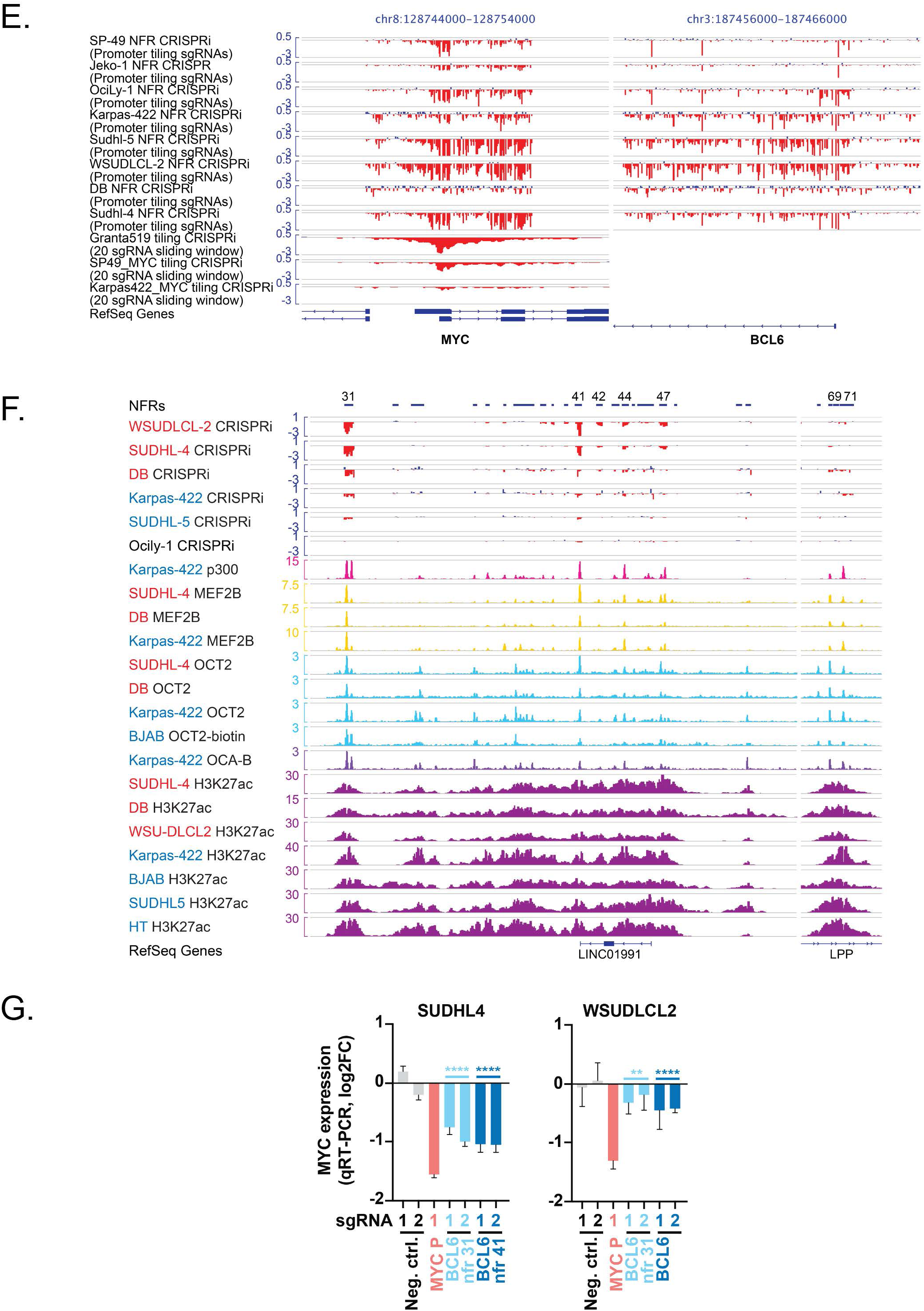
**S3A.** Optical mapping data from three *MYC*::*BCL6*-SE+ lymphoma cell lines showing maps used for parsimonious model of genomic rearrangements (see Figure 3A). **S3B.** Circos plots showing genome-wide rearrangements as identified by optical mapping in three *MYC*::*BCL6*-SE+ lymphoma cell lines. **S3C.** 4C-seq interaction heatmaps for the MYC::BCL6-SE+ cell lines DB (BCL6-SE viewpoint shown) and Sudhl4 (MYC promoter viewpoint show). 4C reads were mapped to a custom genomic map representing a fusion of the MYC and BCL6 loci on the 3’ side of the MYC gene (position shown by dotted line). H3K27ac ChIP-Seq and CRISPRi signal (log2-fold-change) are shown for the corresponding cell lines. **S3D.** Genome browser detail for regions of chromosomes 11 and 1 rearranged to the *MYC* and *BCL6* loci in Sudhl-4 cells, showing H3K27ac ChIP-Seq signal and CRISPRi screen sgRNA depletion and enrichment in the indicated populations. Note lack of essential distal enhancer detection. Note that sgRNAs targeting the promoters of essential genes *TOMM10* and *GGPS1* are depleted in all cell lines regardless of rearrangement. **S3E.** CRISPRi screen sgRNA depletion and enrichment for *MYC* and *BCL6* promoter regions in 8 NFR-focused CRISPRi screens (sgRNAs tiled over both promoters, individual sgRNA depletion shown) and 3 MYC locus tiling sgRNA screens (20 sgRNA sliding window). **S3F.** Genome browser view of the BCL6 super-enhancer region, showing CRISPRi screen sgRNA depletion and enrichment for all 6 GCB-DLBCL screens and ChIP-Seq signal for p300, ternary complex TFs, and H3K27ac in multiple GCB-DLBCL cell lines. *MYC::BCL6*-SE+ cell line names are colored red and GCBME-1-dependent cell line names are colored blue. Genomic coordinates are the same as Figure 3B (hg19 chr3:187610000-187730000 and chr3:187950000-187970000). **S3G.** Change in *MYC* transcript levels (qRT-PCR) in *MYC::BCL6*-SE+ DLBCL cell lines expressing doxycycline-inducible dCas9-KRAB after transduction with indicated sgRNAs targeting *MYC* or *BCL6* locus NFRs (**** p<0.0001, ** p<0.01, t-test of pooled replicates for both sgRNAs targeting the same *BCL6*-SE NFR versus both negative control sgRNAs.

**Supplementary Figure 4.**
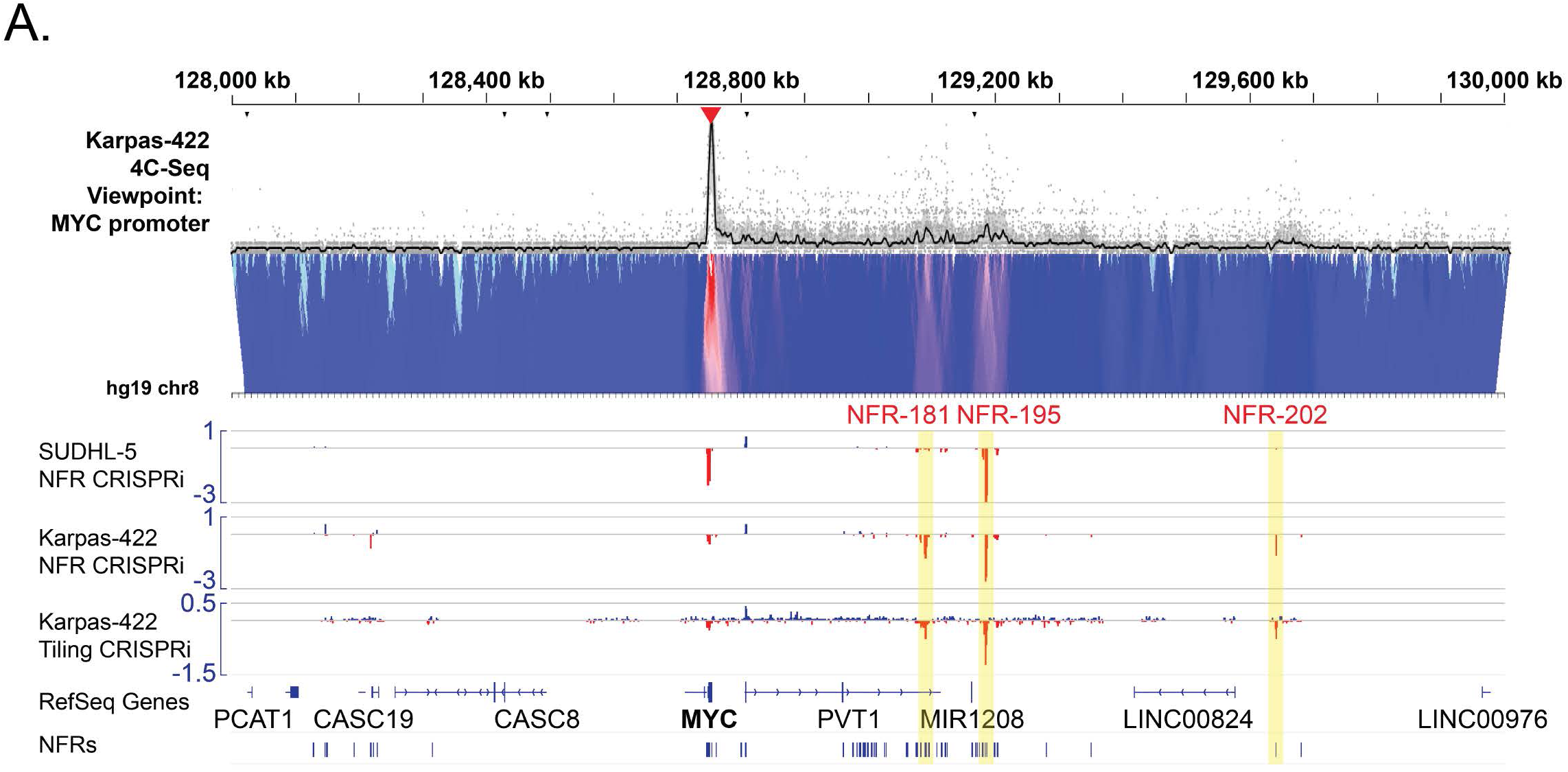
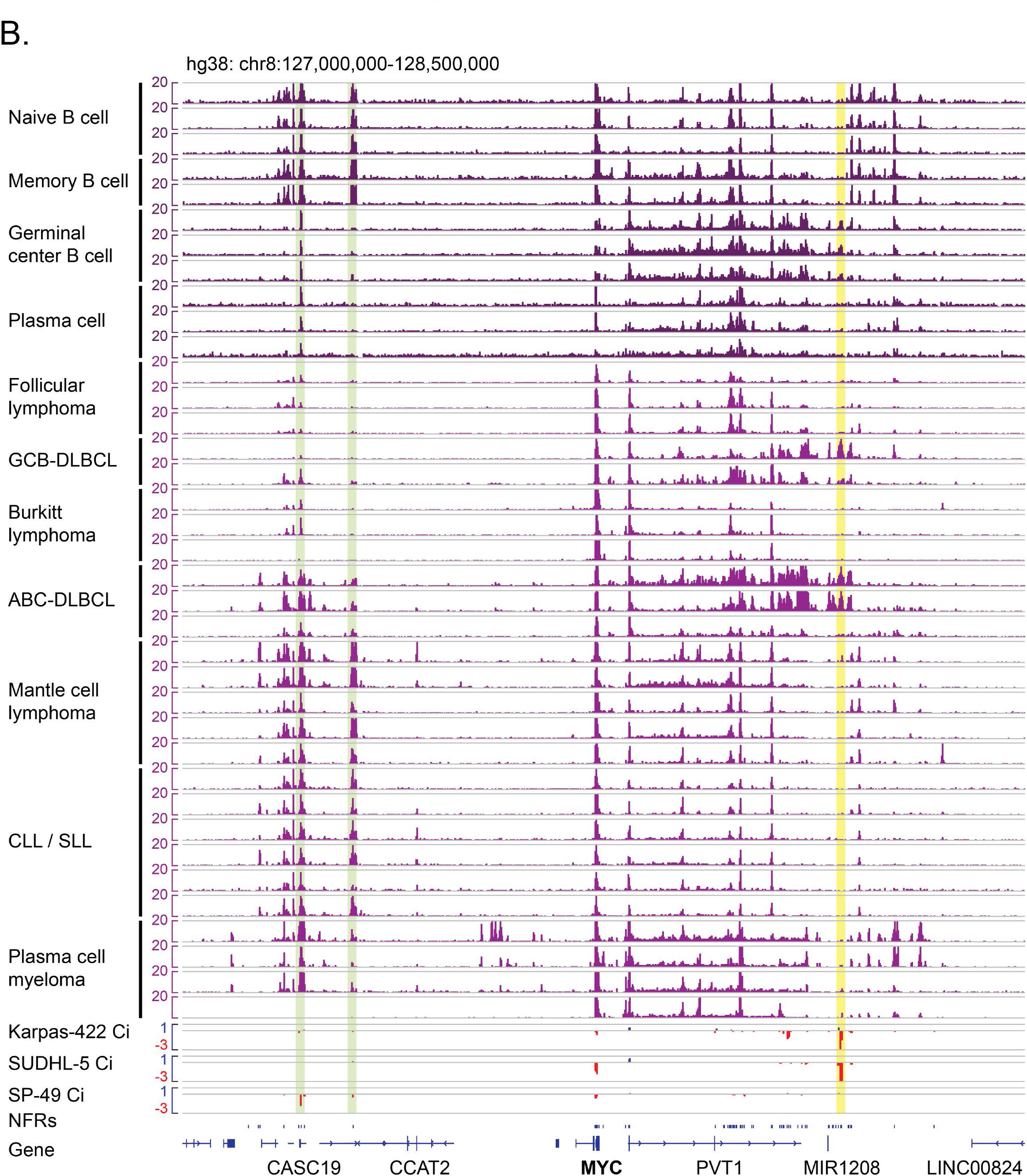
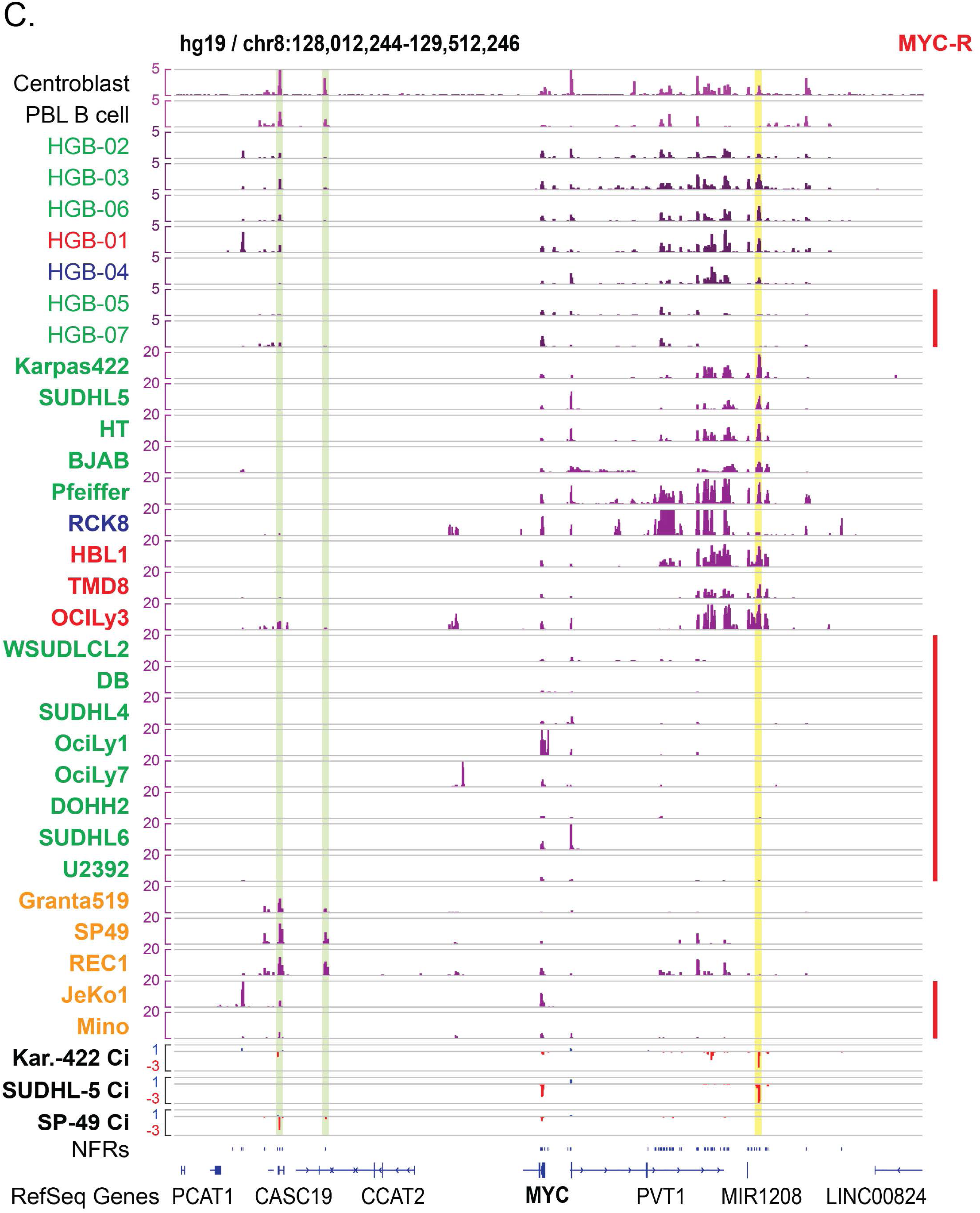
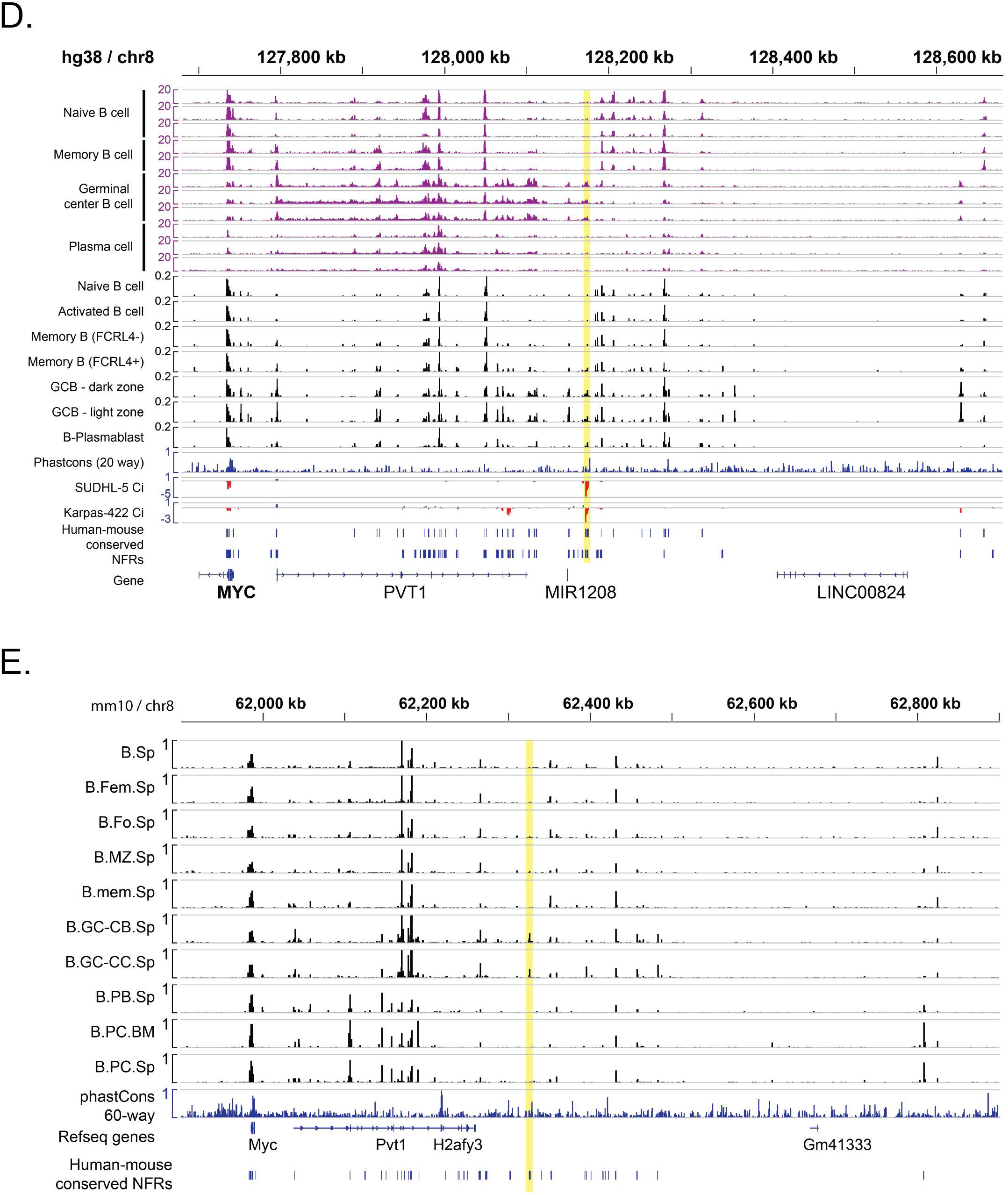
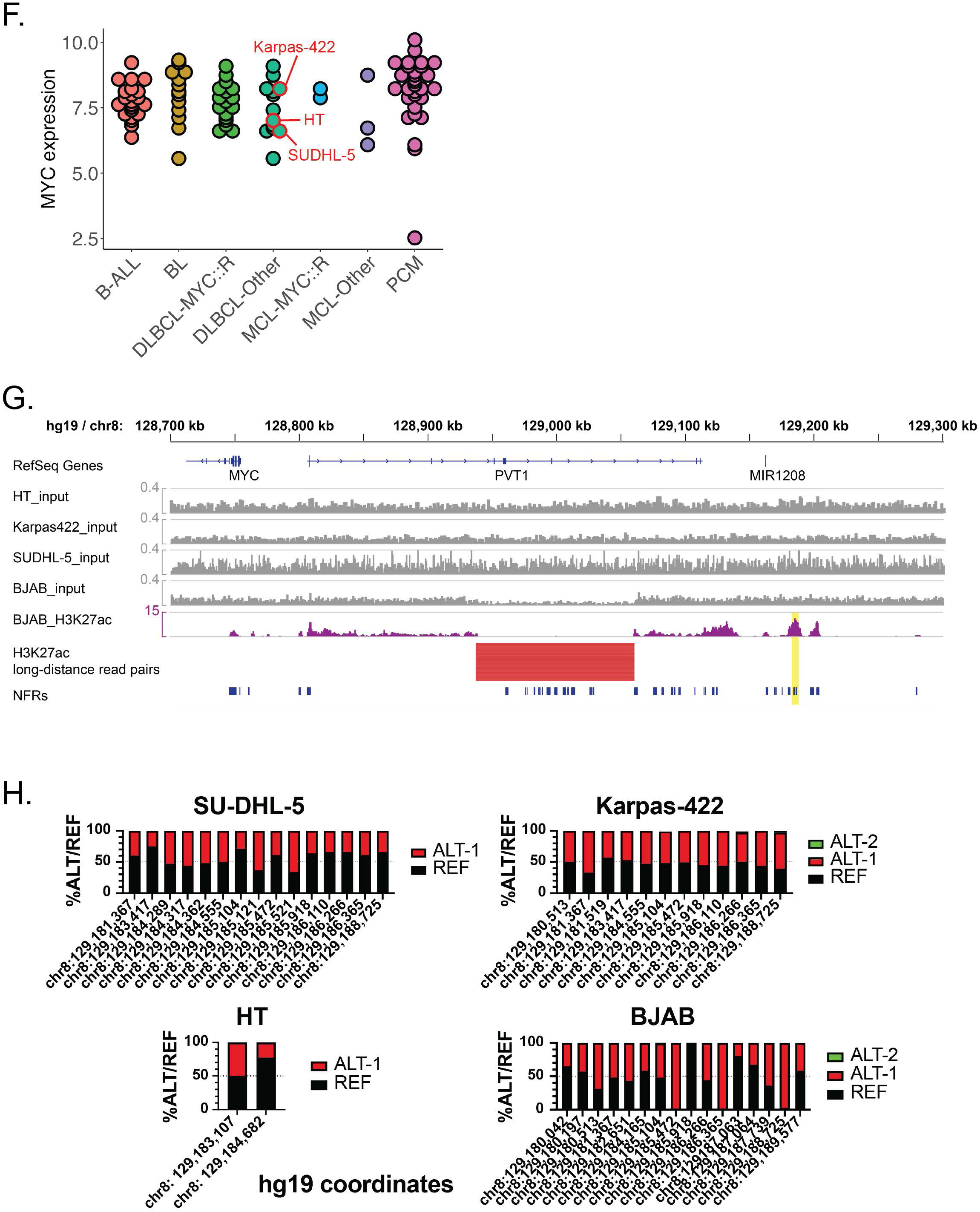
**S4A.** 4C-seq generated in Karpas-422 (4CSeq Pipe analysis) showing genomic interaction frequency with a *MYC* promoter viewpoint. The heatmap at bottom displays median normalized contact intensities at resolutions of 1 kB to 50 kB, while the black trace at top shows median normalized contact intensity at a single resolution. CRISPRi sgRNA depletion / enrichment in SUDHL-5 and Karpas-422 are shown at bottom. **S4B.** H3K27ac ChIP-Seq data across the *MYC* locus in primary normal B cells and mature B-cell neoplasms (Blueprint consortium), showing relationship to essential *MYC* enhancers validated in multiple *MYC*-intact MCL cell lines (green) or multiple *MYC*-intact GCB-DLBCL cell lines (GCBME-1, yellow). **S4C.** H3K27ac ChIP-Seq data across the *MYC* locus in DLBCL and MCL cell lines as well as primary normal B cells and primary DLBCL biopsies (Ryan et al 2015), showing relationship to essential *MYC* enhancers (highlighted as in B). Region is the same as shown in S4Aa. Sample names are color coded as follows: GCB-DLBCL = green, ABC-DLBCL = orange, other / unclassified large B cell lymphoma = blue, MCL = orange. *MYC* rearrangement status is indicated at right. **S4D.** Extended view of *MYC* 3’ interacting regions for normal B cell population H3K27ac ChIP-Seq data (Blueprint consortium) and scATAC-Seq-derived pseudobulk populations (King et al 2021). CRISPRi signal, location of NFRs screened by CRISPRi, and NFRs conserved between human and mouse (see methods) are indicated at bottom. The GCB-ME1 element is highlighted in yellow. **S4E.** ATAC-Seq data for sorted mouse B cell populations (Immgen consortium) across the region of the murine *MYC* locus syntenic to the human region shown in S4C. NFRs conserved between human and mouse are indicated at bottom. The murine element homologous to the human GCB-ME-1 is highlighted in yellow. Note that this element is selectively accessible in murine germinal center centroblasts (“B.GC-CB.Sp”) and germinal center centrocytes (“B.GC-CC.Sp”) **S4F.** *MYC* expression levels in B cell cancer cell lines (DepMap), with DLBCL and MCL cell lines grouped by known *MYC* rearrangement (*MYC*::R) or no demonstrated *MYC* rearrangement (Other). **S4G.** Genome copy number data showing lack of GCBME-1 amplification. H3K27Ac read-pairs show presence of a deletion within the *PVT1* gene body in BJAB. **S4H.** Allelic frequency of sequence variants detected in H3K27ac ChIP-Seq data (>10 total reads) in the acetylated region including GCBME-1 (hg19 / chr8:129,180,000-129,190,000).

**Supplementary Figure 5.**
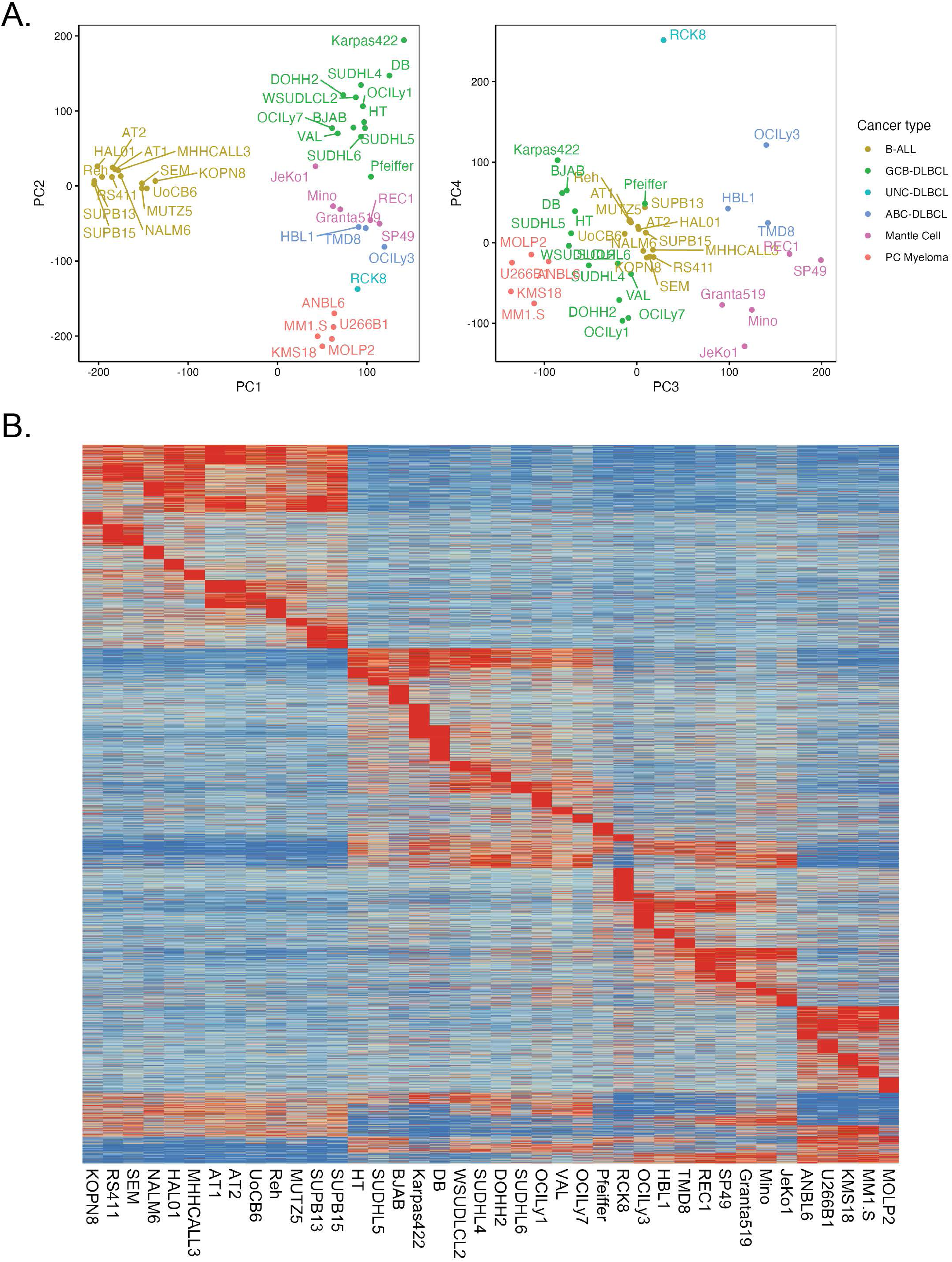
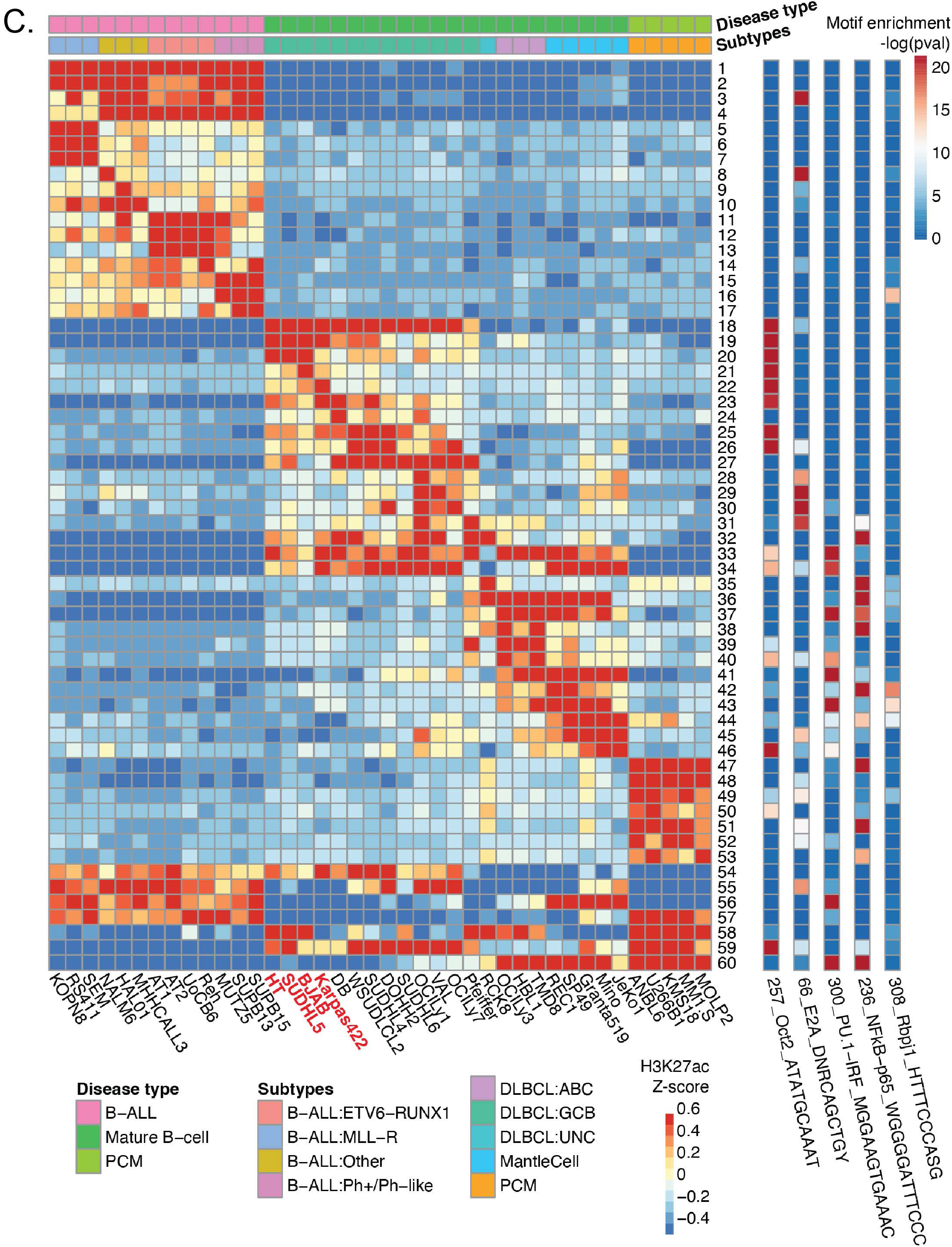
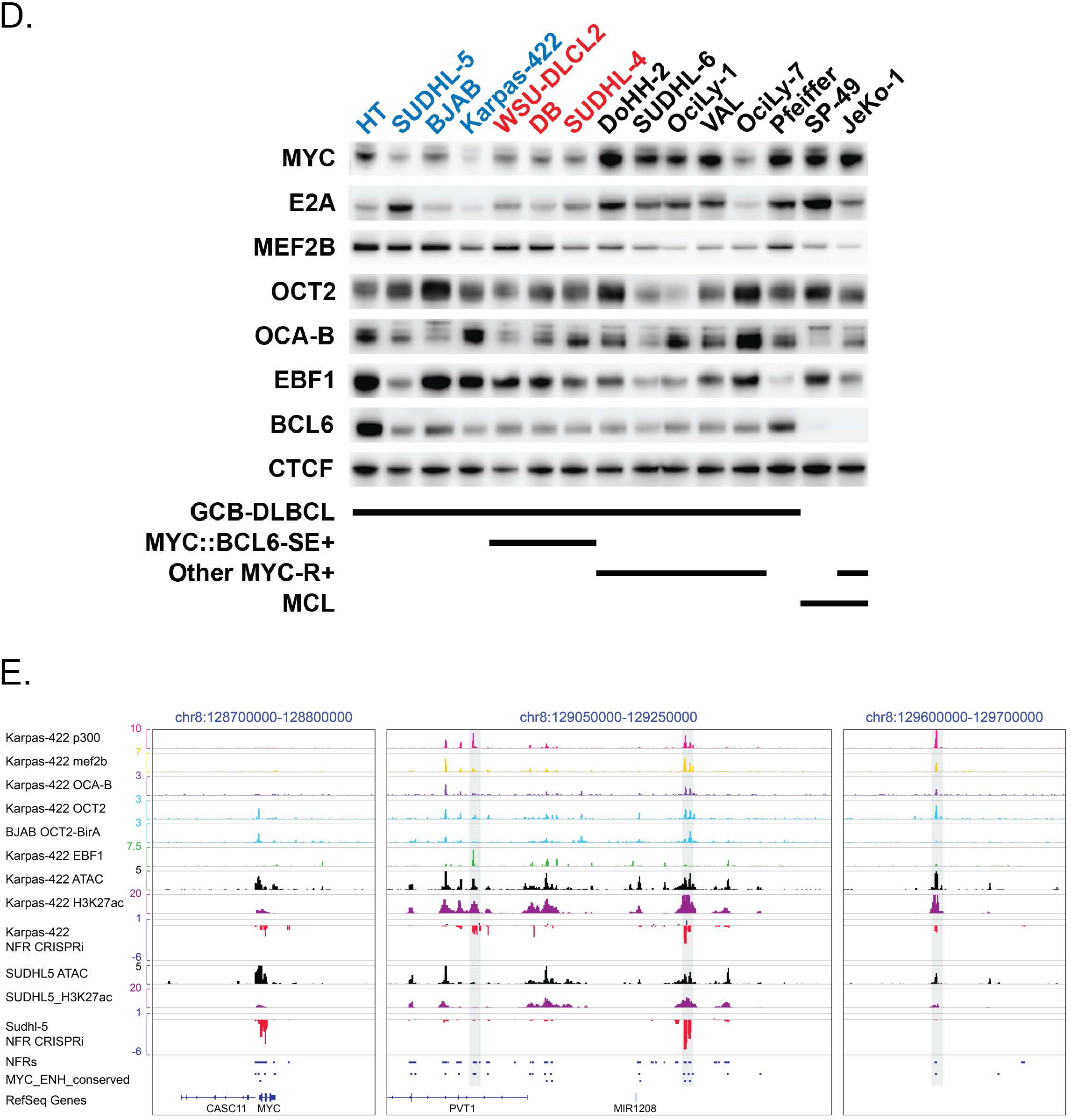
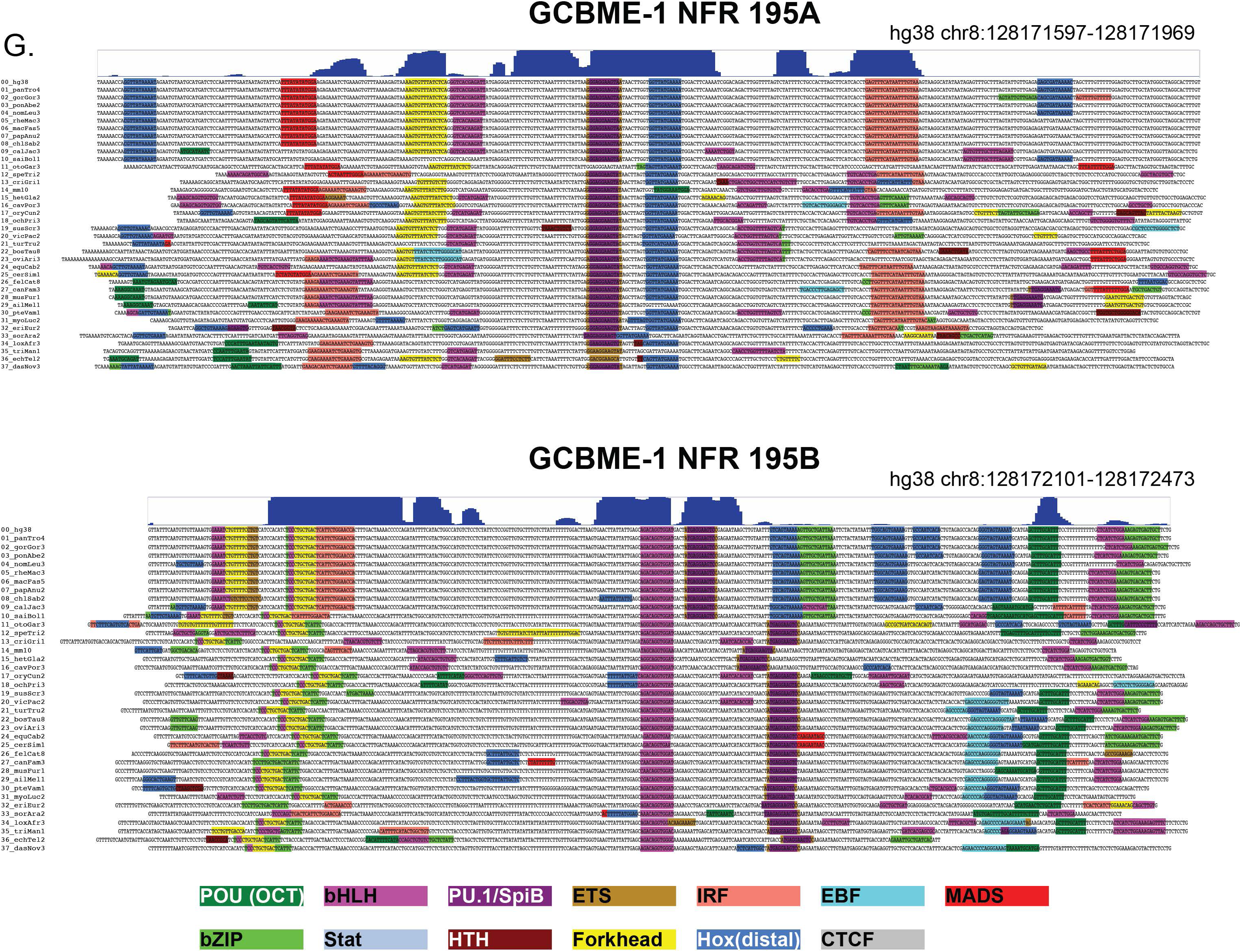
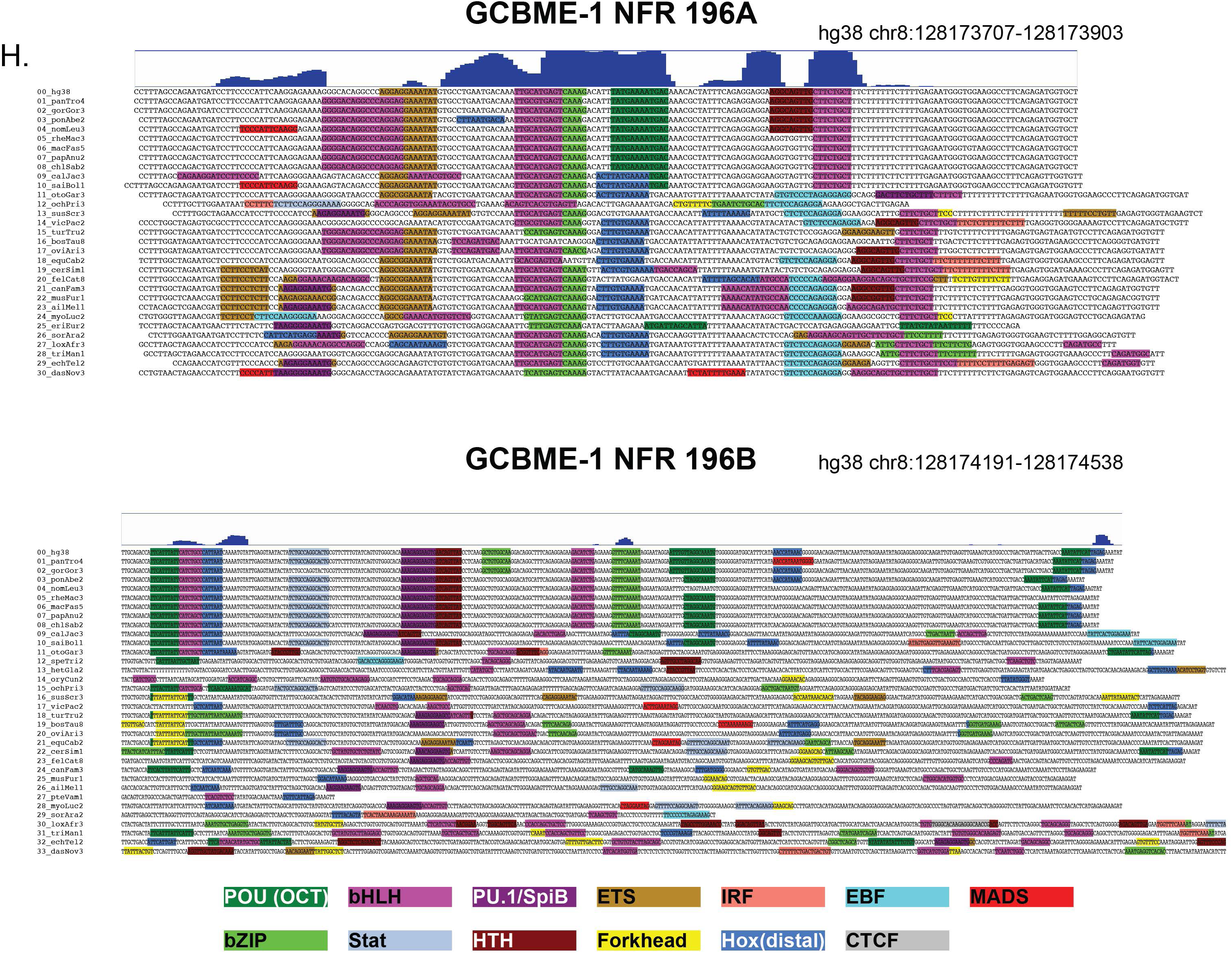
**S5A:** Principal component analysis plots generated from normalized H3K27ac ChIP-Seq signal associated with 98,426 consensus distal NFRs (candidate enhancers) in 40 B cell cancer cell lines. NFRs were defined as consensus ATAC-Seq peaks from datasets generated in the same 40 B cell cancer cell lines. **S5B:** K-means clustering (k=60) of normalized H3K27ac ChIP-Seq signal at 98,426 distal candidate enhancer NFRs / 40 B cell cancer cell lines. **S5C:** Heatmap showing median H3K27ac z-score for each of the 60 enhancer clusters shown in S5B. Cell line cancer type and subtype are indicated at top. Bars at right show results of known motif enrichment analysis (HOMER) for elements in each cluster. Main figure 5A is a subset of data shown here (clusters 18-32, mature B-cell lymphoma cell lines). **S5D:** Western blot of nuclear extracts showing expression of B cell transcription factors in GCBME1-dependent, MYC::BCL6-SE-dependent, and other lymphoma cell lines. MEF2B and CTCF blots shown in Figure 5B are shown again here (with additional cell lines) for comparison. **S5E.** Overview of TF binding, ATAC-Seq, and acetylation for DLBCL enhancer regions identified as essential in Karpas-422 and / or SUDHL5. **S5G.** Motif conservation analysis of NFR 195A and 195B within the GCBME-1. Vertebrate sequences syntenic to the human hg38 region were identified from multiz chain-nets, and ungapped regions were retrieved from the corresponding genomes. HOMER findMotifs was used to identify all above-threshold motifs. Motifs belonging to selected TF binding domain families and individual factors were selected or visualization on the basis of genome-wide enrichment in B cell active enhancers and presence of conserved motifs in the selected sequenced. **S5H.** Motif conservation analysis of NFR 196A and 196B within the GCBME-1, generated as described above.

**Supplementary Figure 6.**
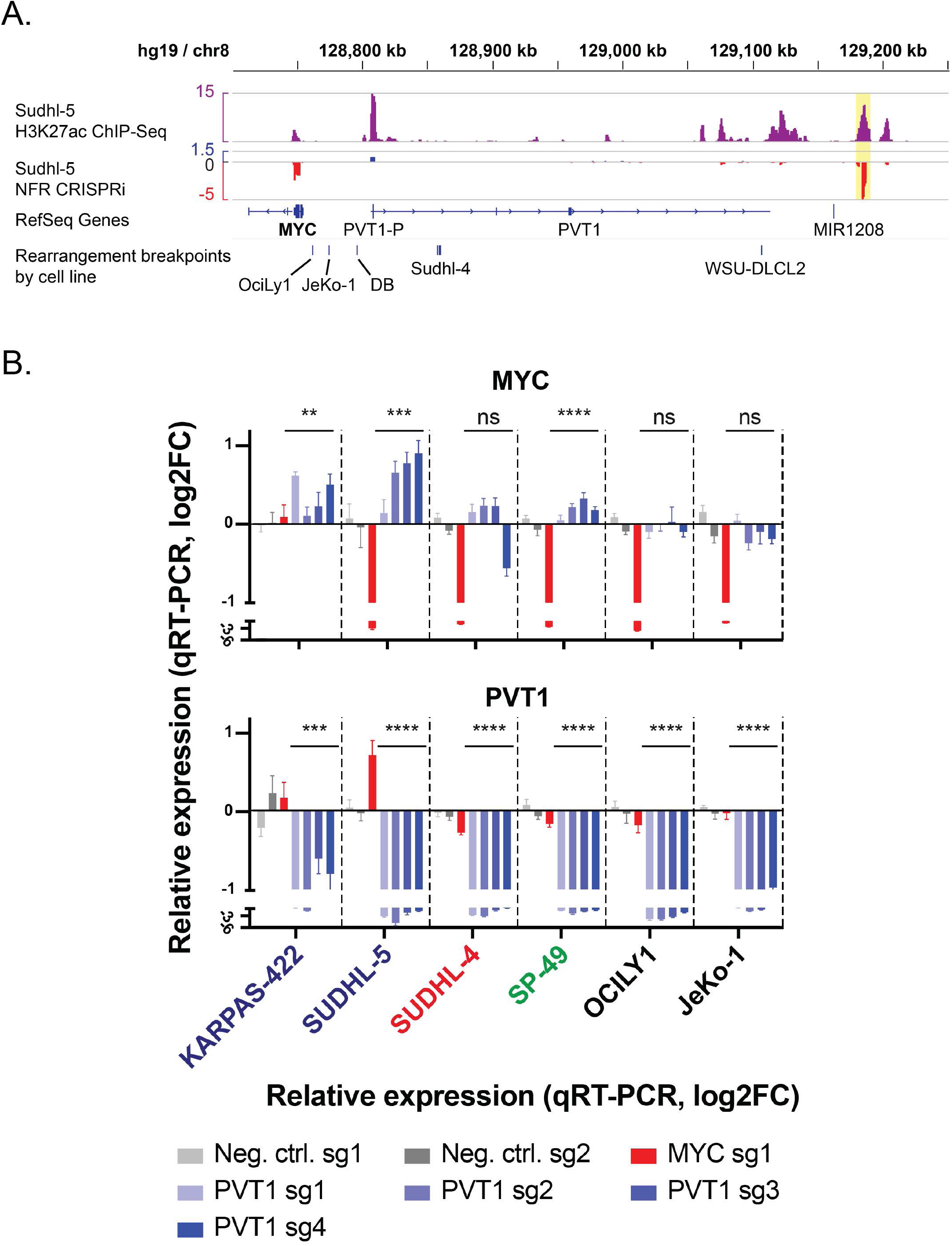
**S6A.** Positions of *MYC* locus breakpoints for the five *MYC*-rearranged lymphoma cell lines screened by CRISPRi in relation to the *PVT1*-P, GCBME-1 (yellow highlight) and essential / acetylated elements in the 3’ *MYC* locus. **B.** Effect of sgRNAs targeting the *PVT1* promoter on *PVT1* and *MYC* transcript levels (qRT-PCR) in lymphoma cell lines bearing doxycycline-inducible KRAB-dCas9. Cell line names are color-coded as follows: Blue, *MYC*-intact GCB-DLBCL; Red, *MYC::BCL6*-SE+ GCB-DLBCL with the breakpoint downstream of the *PVT1* promoter; Green, *MYC*-intact MCL; Black, DLBCL and MCL cell lines with *MYC* rearrangement breakpoint between the *MYC* gene and the *PVT1* promoter.

## Supplementary Tables

Table S1: Oligonucleotide sequences and cell line information

Table S2: Analysis of NFR and gene–focused CRISPRi screens in 8 lymphoma cell lines

Table S3: Analysis of MYC locus tiling CRISPRi screens

Table S4: HOMER motif analysis on enhancer clusters identified in 40 B-cell cancer cell lines

